# Influenza infection recruits distinct waves of regulatory T cells to the lung that limit lung resident IgA+ B cells

**DOI:** 10.1101/2022.09.19.508325

**Authors:** Louisa E. Sjaastad, David L. Owen, Sookyong Joo, Todd P. Knutson, Christine H. O’Connor, Braedan McCluskey, Rebecca S. LaRue, Ryan A. Langlois, Michael A. Farrar

## Abstract

The role of regulatory T cells (Tregs) in limiting responses to pathogens in tissues remains poorly described. We used scRNA-Seq and a newly generated *Foxp3*-lineage reporter line (*Foxp3-iDTR* mice) to track Tregs in the lungs and peripheral blood following infection with influenza virus. Few Tregs of any type were found in the lung at steady-state. Following influenza infection Tregs expressing a strong interferon-stimulated gene signature (ISG-Tregs) appeared by day 3, peaked by day 7, and largely disappeared by day 21 post-infection. A second diverse wave of tissue-repair-like Tregs (TR-Tregs) appeared by day 10 and were maintained through day 21 post-infection. These two distinct Treg subsets had different gene expression patterns and distinct TCR repertoires. To establish the role of Tregs during influenza infection, we acutely ablated Tregs at day 6 post-infection; this resulted in a significant increase in IgA+ B cells in the lung. To determine whether distinct Tregs subsets could also be observed in response to respiratory viral infections in humans we analyzed scRNA-Seq datasets of patients with COVID-19. Peripheral blood from healthy human volunteers had multiple Treg subsets defined by unique gene expression patterns, but few ISG-Tregs. In contrast, two distinct Tregs subsets were expanded in COVID-19 patients - ISG-Tregs and IL32 expressing Tregs (16-fold and 2-fold increased, respectively). ISG-Tregs were present at significantly higher levels in patients with mild versus severe COVID-19, while IL32 expressing Tregs showed the opposite pattern. Thus, the Treg response to respiratory viruses in humans is also diverse and correlates with disease outcome.

## Introduction

Regulatory T-cells (Tregs) protect against autoimmunity and inappropriate immune responses to commensal microorganisms and dampen immune responses to pathogens (Josefowicz et al., 2012; Sjaastad et al., 2021; Veiga-Parga et al., 2013). Although Tregs were initially described as a monomorphic population it soon became evident that they exhibited considerable heterogeneity. Tregs were initially broken down into a population of quiescent central Tregs that circulate through secondary lymphoid tissues and depend on IL-2 for survival, and a second population of effector Tregs that exhibits an activated phenotype, is prevalent within tissues, and requires signaling through ICOS (Smigiel et al., 2014). Subsequent studies demonstrated that during an active immune response, Tregs can express transcription factors that define distinct T-helper cell lineages and enable these Tregs to limit the corresponding T-helper immune response (Harrison et al., 2019; Koch et al., 2009; Sefik et al., 2015; Zheng et al., 2009). Mice bearing Tregs that are deficient in these transcription factors develop uncontrolled inflammation in multiple contexts (Chaudhry et al., 2009; Harrison et al., 2019; Levine et al., 2017; Wohlfert et al., 2011). More recently, single-cell RNA-sequencing (scRNA-seq) studies have allowed the examination of Treg heterogeneity in more depth (Lu et al., 2020; Miragaia et al., 2019; Owen et al., 2022; Owen et al., 2019). These studies demonstrated transcriptional heterogeneity in Tregs within the spleen, lymph nodes, thymus, lung and colon at steady-state, and that Tregs undergo a step-wise acquisition of a tissue-like transcriptome as they move from the lymph nodes to the tissue itself (Delacher et al., 2020; Miragaia et al., 2019). While these studies have profiled Treg diversity at steady state, less is known about the heterogeneity of Tregs within tissues in the context of an activated immune response. In particular, there is limited understanding of how infection impacts Treg heterogeneity within the infected tissue. Herein we demonstrate that Tregs infiltrate the lungs following influenza virus infection and that they do so in distinct waves. An initial wave consisting of Tregs with an IFN-stimulated gene expression signature (ISG-Tregs) and distinct TCR repertoire arises within 7 days post infection before being replaced by a second wave of highly diverse Tregs that express numerous genes associated with tissue repair (TR-Tregs) such as *Areg* and *Tgfb1*. Collectively, Tregs play an important role in limiting the number of IgA+ expressing B cells in the lung. Distinct Treg subsets including the ISG-Treg subset are also observed in human patients following infection with SARS COV2. Healthy volunteers have very few ISG-Tregs but they expand greatly in response to infection with SARS-CoV-2 as does a distinct IL32-expressing subset. Moreover, ISG-Tregs were ∼50% more abundant in patients with mild versus severe COVID-19; conversely, IL32+ Tregs were selectively increased in patients with severe COVID-19. Thus, our findings describe diverse subsets of Tregs that respond rapidly to respiratory viral infections, and which correlate with disease severity in human patients.

## Results

During infection with influenza A virus, Tregs infiltrate and expand within the lung tissue. To examine this in more detail, WT mice were infected with influenza A and analyzed at several time points post-infection. Prior to tissue harvest we injected mice iv with an antibody to CD4 to identify CD4^+^ cells in the blood versus those in the lung tissue (Anderson et al., 2014). In uninfected mice the majority of Tregs isolated from the lungs are in the blood (∼90%) while at 10 days post infection the majority of Tregs are in the lung tissue (∼85%) (Fig. S1A). Over the course of the infection, the number of Tregs within the lungs increased substantially and then contracted following virus clearance (Fig. 1A). This was primarily due to an increased number and frequency of Tregs within the lung tissue (Fig. 1A, 1B). At 10 days post infection, nearly 50% of Tregs within the lung tissue expressed the Th1-associated transcription factor, TBET, while expression of the Th2 and Th17-associated transcription factors, GATA3 and RORүt respectively, remained much lower (Fig. 1C). This result fits with past observations that influenza A virus infection stimulates a strong type I immune response. TBET-expressing Tregs have been identified in several other studies and play an important role in limiting Th1-driven inflammation. While this result demonstrated that TBET+ Tregs expand in the lungs during influenza A infection, the degree of Treg heterogeneity during an influenza infection and their functions within the lungs during the infection remained unclear.

**Figure 1.**
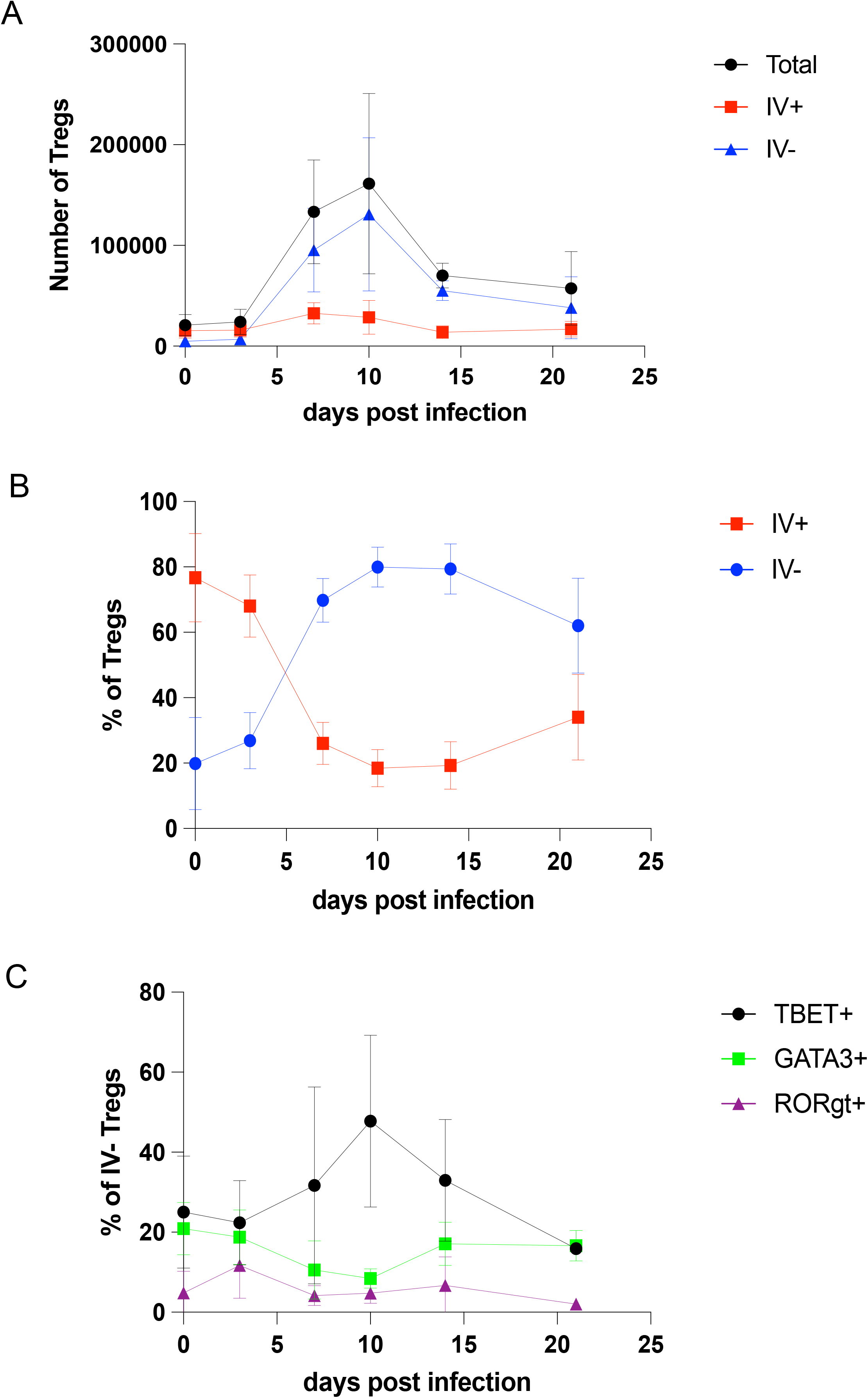
Tregs expand in the lungs during influenza A virus infection. (**A**) Number of Tregs within the lungs and divided between the IV+ and IV- fractions over the course of influenza A infection. (**B**) Frequency of Tregs within the IV+ and IV- fractions of the lungs over the course of influenza A infection. (**C**) Frequency of Tregs within the IV- fraction of the lungs expressing TBET, RORүt, and GATA3. N=4-8 per time point. Horizontal bars represent means and vertical bars represent standard deviations.

To interrogate the phenotype of Tregs responding to the lungs during influenza infection, we performed single-cell RNA-sequencing (scRNA-seq). *Tclib+ x Tcrva*^*+/-*^ *x Foxp3-GFP* mice received temporally staggered infections with influenza A virus (PR8 strain)(Fig. 2A). Prior to tissue harvest we injected mice iv with a CITE-Seq antibody to CD4 to identify CD4^+^ cells in the blood (Ab^+^) versus those in the lung (Ab^−^) (Anderson et al., 2014). We used oligonucleotide-conjugated tags (“hashtag” antibodies) to identify cells obtained at different timepoints after infection, as well as mark distinct biological replicates. CD4^+^FOXP3^+^ Tregs were isolated from the lung by FACS and single cell libraries were generated. After quality control and removal of doublets, transcriptomes from 17,938 combined single cells were analyzed. A dimensional reduction with graph-based clustering approach was applied to combined data from all five timepoints to identify distinct Treg subsets; this resulted in a final UMAP plot with thirteen distinct clusters (Fig. 2B, Table S1. Tregs derived from experimental replicates at each time point were phenotypically reproducible (Fig. S1B). Distinct Treg subsets were present within the lung tissue and blood fractions and arose with different kinetics (Fig. 2B, 2C, S1C). A distinct cluster of ISG-Tregs (cluster 3) emerged at 7 days post infection (dpi) and contracted upon resolution of the infection (10 and 21 dpi) (Fig. S1C, S1D, 2C). In the ISG-cluster, all the top differentially expressed genes were ISGs. In addition to the ISG-Treg cluster, this analysis also demonstrated intriguing dynamic heterogeneity within the lung Tregs across the infection time course. Cluster 1 Tregs were present primarily in the blood and expressed transcripts found in central Tregs, such as *Ccr7* (Fig. S1D). Cluster 4 Tregs appear to represent a transitional early effector Treg subset that branches off into three distinct lineages (Fig. 2D) that we will tentatively refer to as tissue-resident-like Tregs (cluster 0), tissue-repair-like (TR)-Tregs (clusters 2, 5, 6, 8, 10, and 12) and ISG-Tregs. Cluster 0 was defined by transcripts indicative of early effector-like Tregs such as *S100a10, S100a4, and S100a6* (Fig. S1D) (Permanyer et al., 2021; Zemmour et al., 2018). Transcripts for *Areg* were highly expressed in cluster 2, indicative of Tregs with tissue-repair functions (Fig. S1D). This is consistent with studies demonstrating that deletion of *Areg* in all Tregs leads to substantial defects in lung function in mice infected with influenza (Arpaia et al., 2015). Cluster 2 was closely linked to clusters 5, 6, 8, 10 and 12, which we collectively refer to as TR-Tregs. Tregs within cluster 5 express gene transcripts associated with activation including *Ccl5* and *Nkg7* (Fig. S1D). Cluster 6 represents granzyme B-expressing Tregs (Fig. S1D), which have a previously described role in limiting lung inflammation during RSV viral infection (Loebbermann et al., 2012). Cluster 8 Tregs uniquely expressed transcripts for *Plac8* (Fig. S1D), a placental protein that has been observed in Tregs but whose function is unclear. Cluster 10 Tregs expressed transcripts for *Penk* (Fig. S1D), an endogenous opioid, that was recently shown to be expressed in Tregs from UVB-irradiated skin that promote wound healing (Shime et al., 2020). Finally, Tregs within cluster 12 expressed *Tff1* (Fig. S1D), a member of the trefoil factor protein family, which has been shown to promote repair of gut epithelium (Playford et al., 1996). Transcripts for *Tff1* have been found in IL18R^+^ Tregs within the thymus (Peligero-Cruz et al., 2020) while accessible chromatin at the *Tff1* locus has been identified in ST2-expressing Tregs from the colon and skin (Delacher et al., 2020). Thus, Tregs in clusters 2, 5, 6, 8, 10, and 12 all express transcripts that have been associated with distinct aspects of tissue-repair.

**Figure 2.**
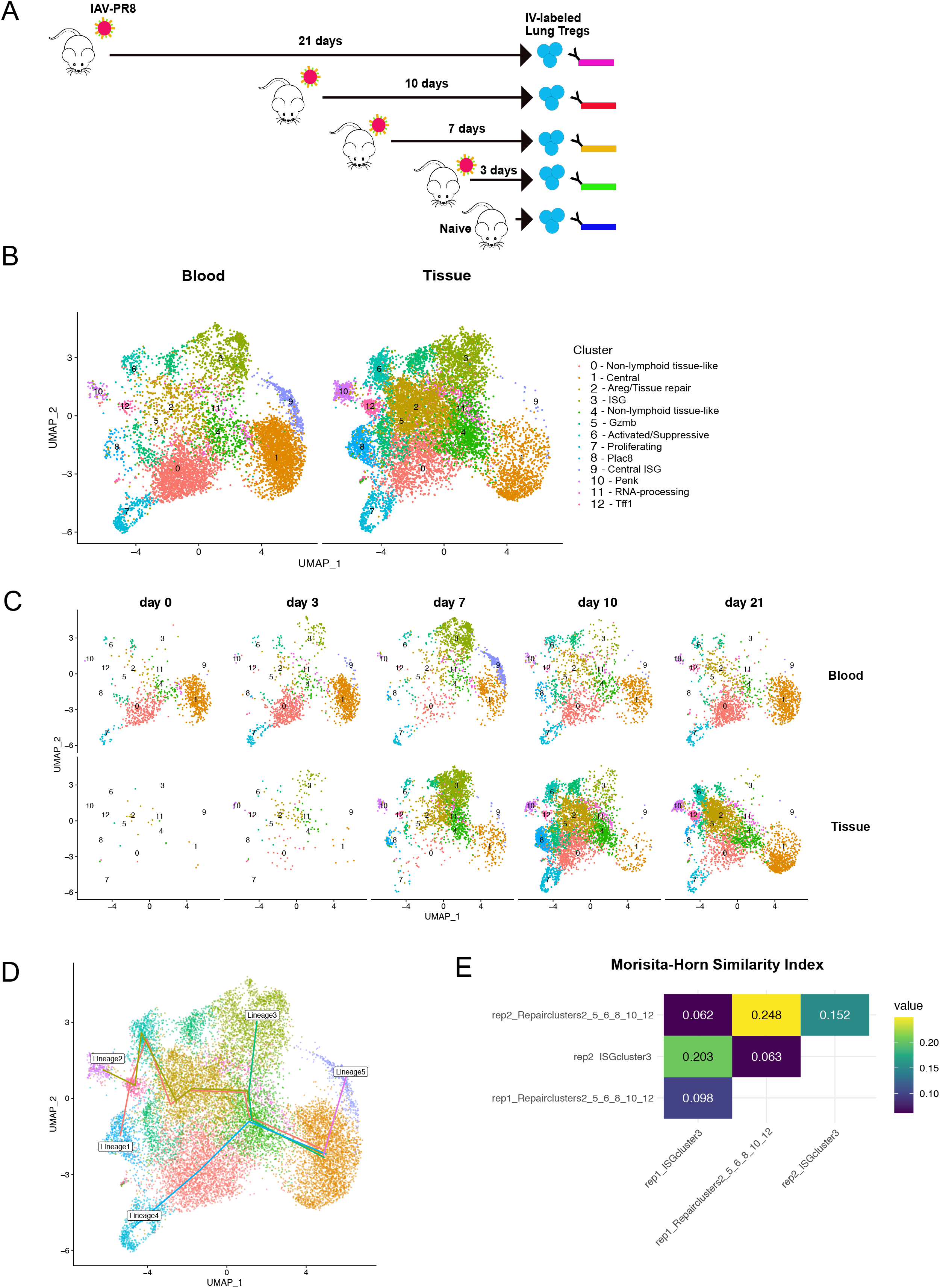
Treg heterogeneity during influenza A virus infection. (**A**) Experimental scheme for single-cell RNA-seq of lung Tregs from influenza A virus infected mice. Infections were staggered and IV-labeled Tregs were sorted from the lung and hashtagged prior to sequencing. (**B**) UMAP of Tregs clustered at resolution 0.5 and separated by IV-labeling status. (**C**) UMAP of Tregs separated by days post infection and IV-labeling status. (**D**) Slingshot trajectory analysis. Morisita-horn similarity index of Treg TCRs among ISG-Tregs (cluster 3) and TR-Tregs (clusters 2, 5, 6, 8, 10, and 12) from individual mice.

ISG-Tregs are distinct from other Treg lineages that migrate to the lung during influenza virus infection. Using slingshot trajectory analysis, we observed that central Tregs (cluster 1) progress into cluster 4 Tregs (an early effector Treg-state) and then branch off into 3 distinct lineages: ISG-Tregs, TR-Tregs (clusters 2, 5, 6, 8, 10, 12), and tissue-resident-like Tregs (cluster 0) (Fig. 2D). In support of this conclusion, TCR repertoire analysis revealed a high degree of similarity between Treg TCRs from biological replicate mice within the ISG-Treg cluster (Fig. 2E). Likewise, there was considerable overlap between TCRs from TR-Tregs between independent mice. In contrast, much lower similarity between TCRs was observed when comparing ISG-Tregs with the TR-Treg subset, even when these populations came from the same mouse (Fig. 2E). This exclusive relationship held even when comparing ISG-Treg TCRs with TCRs from Tregs in each of the other clusters alone (Fig. S2A). Finally, TR-Treg clusters expressed *Itgav* and *Itgb8*, which pair to form integrin αvβ8 (Fig. S2B). These Tregs also express *Tgfb1* mRNA (Fig. S2B). Previous studies have shown that integrin αvβ8 on Tregs is important for the cleavage and activation of TGFβ and subsequent suppression of effector T-cell responses during inflammation(Worthington et al., 2015). Thus, TR-Tregs may be involved in lung remodeling following influenza virus infection. This pattern of transcript expression was not characteristic of all integrins, as transcripts for integrin β1 were preferentially expressed in cluster 0 Tregs (Fig. S2B). TR-Treg clusters also expressed higher levels of transcripts for chemokine receptors, *Cxcr6* and *Ccr8*, as well as activation induced markers, *Klrg1, Il1r2*, and *Pdcd1* (Fig. S2B). In contrast, ISG-Tregs had limited expression of markers characteristic of TR-Tregs, but instead uniquely expressed high amounts of the chemokine *Cxcl10* (Fig. S2B). Thus, ISG-Tregs are a unique Treg lineage that appear early in the lung tissue during influenza virus infection and are phenotypically distinct from other Treg lineages that arrive later.

We have not found reliable surface markers to specifically identify ISG-Tregs. Therefore, to interrogate the dynamics and function of ISG-Tregs, we designed a mouse model that allows one to track distinct Treg lineages using CRE-reporter mice. Specifically, we used CRISPR/CAS9 based approaches to insert a *LoxP*-flanked *STOP* cassette upstream of the *IRES-DTR-GFP* sequence of *Foxp3-DTR* mice (Kim et al., 2007) to enable CRE-inducible expression of DTR and GFP (Fig. S3A). Insertion of the floxed STOP cassette was confirmed by PCR and whole genome sequencing (Fig. S3B, S3C). In the absence of CRE the DTR-GFP cassette is not expressed. In contrast, *Cd4-Cre x Foxp3-iDTR/WT* heterozygous mice resulted in ∼50% of splenic Tregs (CD4^+^CD25^+^ T-cells) expressing GFP (Fig. S3D). *Foxp3-iDTR* mice also allow for inducible expression of DTR/GFP when crossed with mice expressing a tamoxifen-inducible CRE such as CD4-CRE^ERT2^ (Fig. S3E). Finally, we confirmed that diphtheria toxin (DT) administration deletes labeled GFP^+^ Tregs in *Cd4-Cre*^*ERT2*^ *x Foxp3-iDTR* mice just as efficiently as in the original *Foxp3-DTR* mouse (Fig. S3F). To identify ISG-Tregs we crossed *Foxp3-iDTR* mice to *Mx1-Cre* mice, as *Mx1* is a well-characterized interferon-stimulated gene (Kühn et al., 1995). At steady state, 6.8± 2.5% of Tregs within thymus, and 4.0±0.4% of Tregs in spleen and mesenteric lymph nodes expressed GFP (Fig. S3G). The frequency of ISG-Tregs identified in the Mx1-Cre x Foxp3-iDTR mice was similar to that which we observed when reanalyzing a previous scRNA-seq dataset for thymic Tregs and Treg progenitors (Owen et al., 2019), in which ISG-Tregs comprised ∼8% of thymic Tregs and Treg progenitors. Subsequent scRNA-seq studies found a similar population of ISG-Tregs in the spleen (∼5%) (data not shown). This population of Tregs is similar to one identified in the spleen, lung, gut, and skin at steady state and following treatment with IL-2 mutein (Lu et al., 2020; Miragaia et al., 2019). Thus, the percentage of Tregs labeled in *Mx1-Cre x Foxp3-iDTR* mice closely mimics that detected in the spleen and thymus by scRNA-seq.

To determine the origin and fate of ISG-Tregs during influenza infection, we utilized the *Mx1-Cre x Foxp3-iDTR* mouse model. Single-cell RNA-seq data demonstrated that ISG-Tregs peak in the lung 7 days post-infection and then contract upon clearance of the infection (Fig. 2C). However, it is unclear whether ISG-Tregs give rise to the subsequent TR-Treg lineage or not. In *Mx1-Cre x Foxp3-iDTR* mice, ISG-Tregs are permanently labeled with GFP and DTR. In naïve mice, GFP^+^ ISG-Tregs represent ∼4.0% of Tregs within the lung tissue; this expanded to ∼22.1% GFP^+^ at 7 days post infection (Fig. 3A, 3B). While the frequency of GFP^+^ Tregs remained high at 10 days post infection, this decreased precipitously by 21 days post infection (Fig. 3B). A similar pattern of expansion and contraction of GFP^+^ ISG-Tregs occurred in the mediastinal LN (Fig. 3B). Thus, the second wave of Tregs in the lung are not derived from ISG-Tregs.

**Figure 3.**
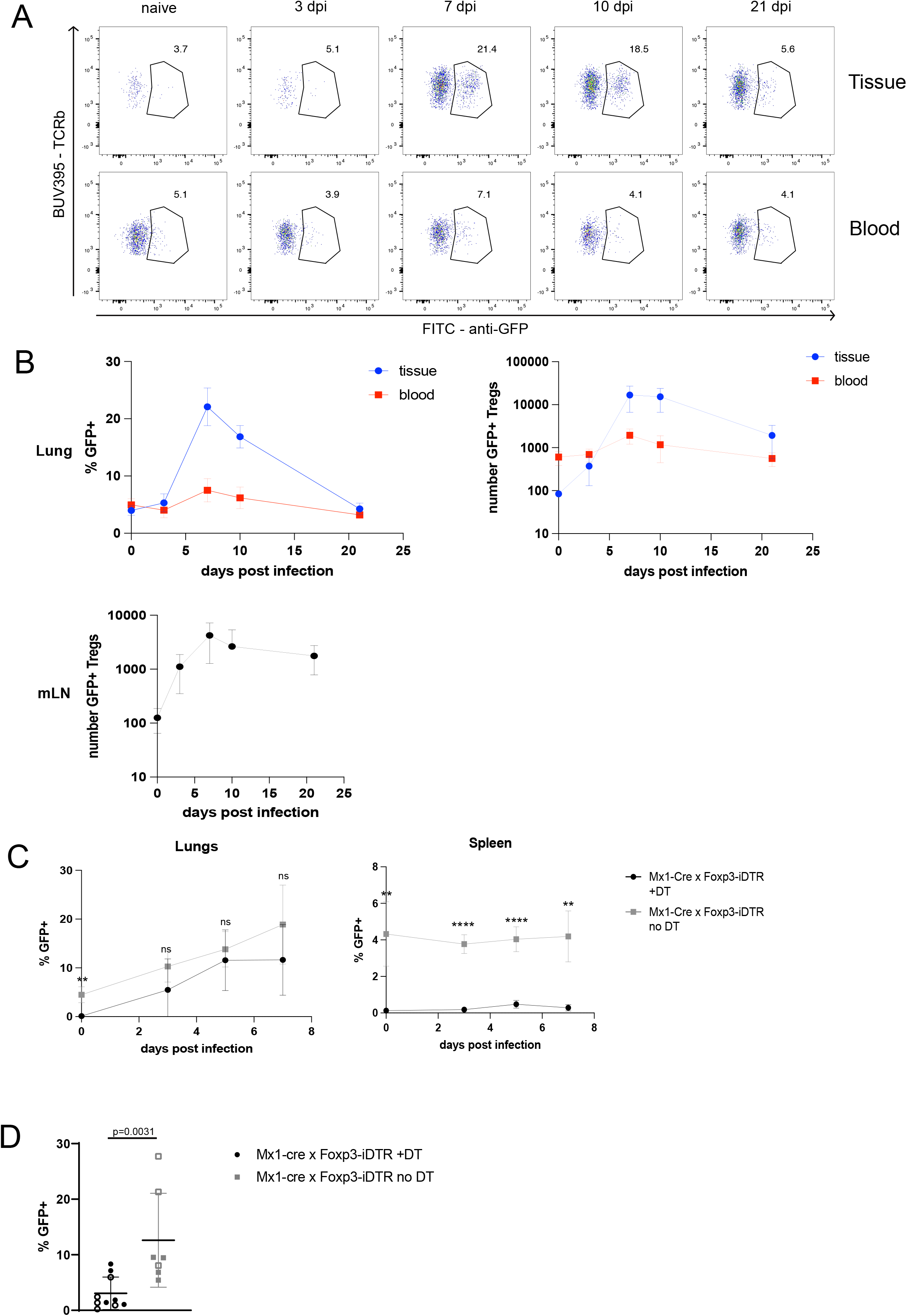
Tracking ISG-Tregs using *Mx1-Cre x Foxp3-iDTR* mice. (**A**) ISG-Tregs in the lungs of *Mx1-Cre x Foxp3-iDTR* mice separated by time point post infection and IV-status. (**B**) Frequency and numbers of ISG-Tregs in the lungs and mediastinal LN of *Mx1-Cre x Foxp3-iDTR* mice across time course. For the lungs, n=5 for uninfected and n=6 for all other time points. For mediastinal LN n=3 for uninfected and n=6 for all other time points. Horizontal bars represent means and vertical bars represent standard deviations, samples are from 2-independent experiments. **(C)** Frequency of ISG-Tregs in the lungs and spleen following pre-infection depletion of ISG-Tregs. Data is representative of 2 experiments, n=3-4 mice per time point. Horizontal bars represent the means and vertical bars represent the standard deviations. **P<0.01, ****P<0.0001, determined by unpaired t-test with Holm-Sidak multiple comparisons correction. (**D**) Frequency of GFP+ Tregs within the lung 7 days following secondary influenza infection. Mice were infected with 1000pfu X31 and rechallenged with 15pfu of PR8 after 28 days. Open and closed symbols represent samples from 2 separate experiments. For Mx1-Cre x Foxp3-iDTR +DT n=10, and for Mx1-Cre x Foxp3-iDTR no DT n=7. Horizontal bars represent means and vertical bars represent standard deviations. P-value determined by Mann-Whitney test.

Since a subset of ISG-Tregs exists in the spleen and lymph nodes of naïve mice, we examined whether ISG-Tregs in the lung following influenza infection arise exclusively from this pre-existing subset, or whether they can be induced to adopt this phenotype. Two days prior to infection we depleted ISG-Tregs with DT. Depletion in the lungs and spleen was quite effective (Fig. 3C). ISG-Tregs in the spleen were depleted for the entire time course of infection (Fig. 3C). In contrast, although ISG-Tregs were effectively depleted in the lungs at day 0, they had fully recovered by day 3 post-infection and remained at equivalent levels to that seen in undepleted mice out to at least day 7 post-infection (Fig. 3C). It is unclear whether this reflects rapid expansion of a small set of pre-existing ISG-Tregs that were not effectively depleted or whether non ISG-Tregs are recruited to the lung upon deletion of the pre-existing ISG-Tregs, and these Tregs then acquire an ISG-signature. As ISG-Tregs contract in the lungs following clearance of influenza, we assessed whether they could be recalled during a secondary infection, and if this too was independent on the pre-existing population or merely reflected IFN abundance in the infected tissue. To examine this issue *Mx1-Cre x Foxp3-iDTR* mice were given a primary infection with influenza X31. The ISG-Tregs were then depleted at 28 days post infection via DT administration. One day after ISG-Treg depletion the mice were rechallenged with influenza PR8 and then analyzed 7 days following the secondary infection. The frequency of ISG-Tregs during the secondary infection was significantly diminished in mice depleted of ISG-Tregs prior to infection compared to control mice (Fig. 3D). This finding demonstrates that ISG-Tregs seen in a secondary infection are dependent on the pre-existing ISG-Treg subset and do not simply arise because they migrate to a tissue with significant IFN production.

To understand the function of Tregs during influenza A infection, we analyzed whole lung transcriptomes of Treg-depleted mice. *Mx1-Cre x Foxp3-iDTR* and *Foxp3-DTR* mice were infected with influenza A virus and depleted of ISG- or total Tregs, respectively, at days 6, 7, and 8 post infection. RNA-seq of whole lung demonstrated that depletion of ISG-Tregs in *Mx1-Cre x Foxp3-iDTR* mice had minimal effect on gene transcription (data not shown). This may reflect our inability to effectively deplete this population in the lungs. However, analysis of Treg-depleted *Foxp3-DTR* mice revealed an increase in transcripts for immunoglobulin genes (Fig. 4A). In particular, transcripts for *Iga* and *Igj* (J-chain) were highly upregulated in the absence of Tregs in depleted *Foxp3-DTR* mice (Fig. 4A). To validate this result, we evaluated the lungs of Treg-depleted *Foxp3-DTR* mice by flow cytometry and found an increased frequency of IgA-expressing B-cells (Fig. 4B). Previous studies have supported a protective role for secreted IgA during influenza infection (van Riet et al., 2012). Our results suggest that Tregs present during peak anti-viral immune activity against influenza A virus serve to limit IgA production by B-cells within the lungs.

**Figure 4.**
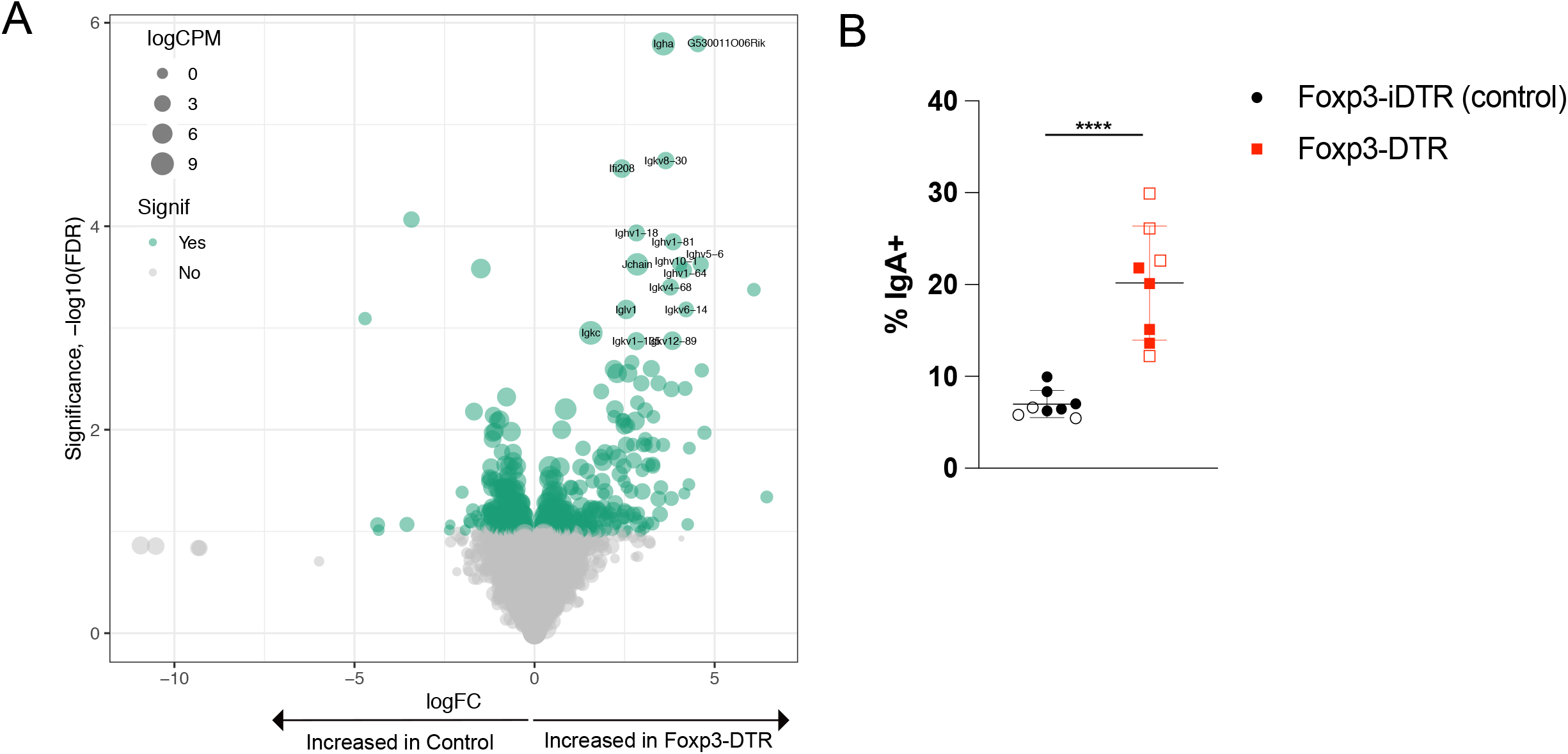
Increased IgA+ cells in the lungs of influenza A infected mice following depletion of Tregs. **(A)** Volcano plot showing changes in gene expression between Treg depleted *Foxp3-DTR* mice and control mice. (**B**) Frequency of IgA^+^ B-cells following depletion of Tregs in *Foxp3-DTR* mice. Gated on live>CD45+>Thy1.2->B220+/IgG H+L+>GL7-cells. N=8 for each treatment group. Open and closed symbols represent samples from 2 separate experiments. Horizontal bars represents means and vertical bars represents standard deviations. ****p<0.0001, determined by student’s t-test.

The role of IFNs in the severity of COVID-19 is controversial with some studies supporting a protective role and others implicating them in progression to severe disease (Lee and Shin, 2020). To examine this in more detail and to determine whether ISG-Tregs exist in humans with respiratory viral infections and correlate with COVID-19 severity, we reanalyzed a scRNA-seq meta-analysis of PBMCs obtained from 111 healthy donors and patients with mild or severe COVID-19 (Mukund et al., 2021). We extracted data for CD4^+^ and CD8^+^ T cells and identified cluster 9 as Tregs based on high expression of *FOXP3* and *IL2RA* transcripts but low expression for *IL7RA* (Fig. S4A). Upon reclustering the Treg subset, we identified 6 distinct clusters; cluster 4 expressed an IFN gene signature similar to that observed in mice (Fig. 5A, Table S2). Additional clusters included ones characterized by the inflammatory cytokine *IL32* (cluster 0), the gene *S100A9* (cluster 1), an activated subset expressing immediate early genes such as *FOS* and *JUN* (cluster 2), a subset expressing of *TCF7* and *PLAC8* (cluster 3), and a subset expressing *GZMA, GZMK, CCL5* and *NKG7* (cluster 5). We separated Tregs based on disease status of the patients from whom they were collected; we found that healthy donors had a very small population of ISG-Tregs. In contrast, patients with both mild and severe COVID-19 exhibited a significant increase in ISG-Tregs (Fig. 5B). Notably, ISG-Tregs were >50% more abundant in patients with mild COVID-19 as opposed to those with severe disease (1.6-fold↑) (Fig. 5B, 5C). There was no significant difference in the time after disease onset when samples were collected (Fig. S4B). The opposite relationship was observed for the *IL32*-expressing Treg cluster, as patients with severe COVID-19 showed a significant increase in *IL32*-expressing Tregs relative to healthy donors (2.9-fold↑) or patients with mild disease (1.9-fold↑) (Fig. 5B, 5C). Thus, ISG-Tregs correlate with better outcomes while IL32-Tregs correlate with worse outcomes in COVID-19 patients.

**Figure 5.**
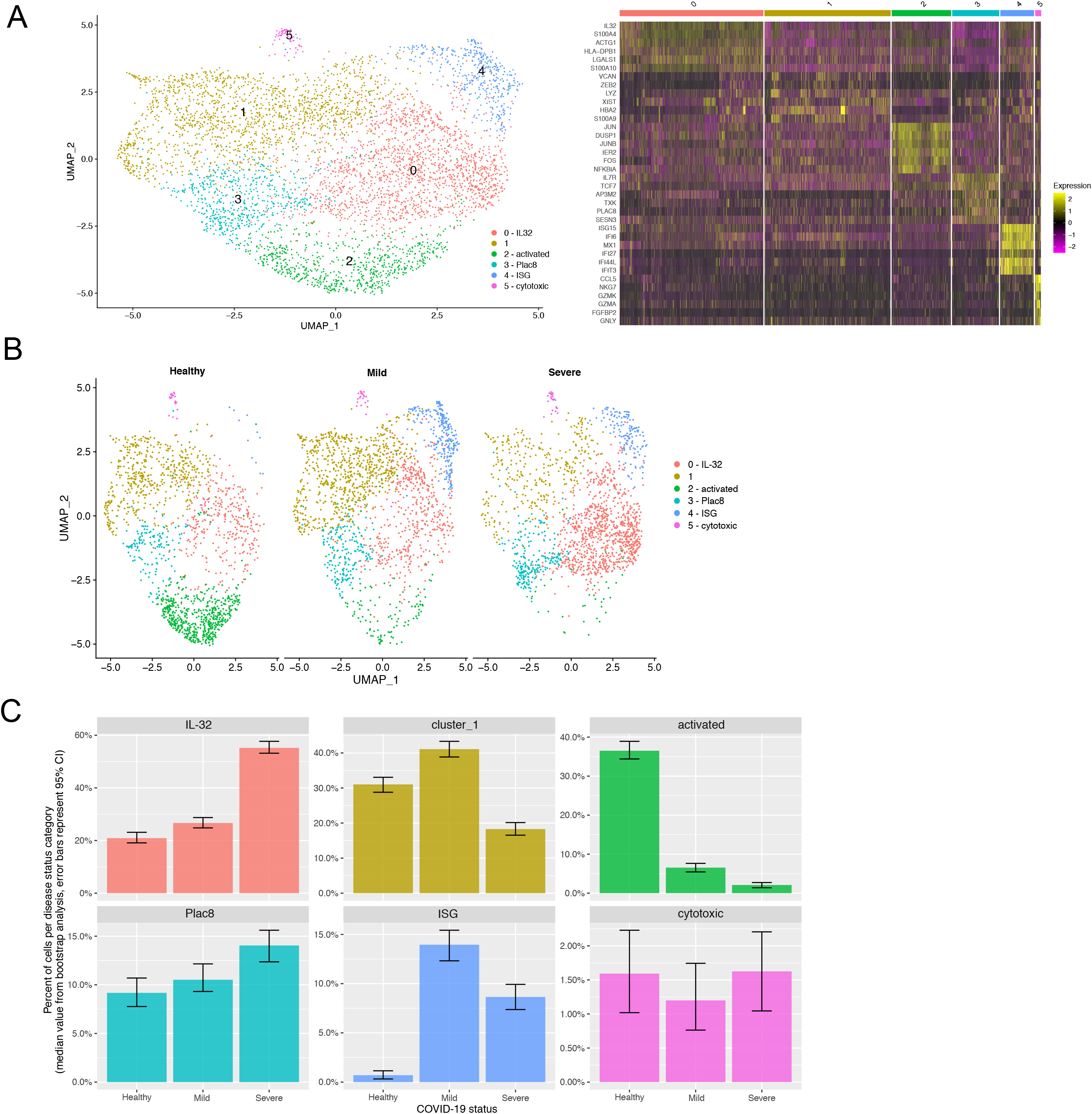
ISG-Tregs in COVID-19. (**A**) UMAP of Tregs from the blood of healthy donors and COVID-19 patients (resolution 0.3). Heatmap of top differentially-expressed genes in each cluster. (**B**) UMAP of Tregs separated by disease status. (**C**) Proportion of Tregs within each cluster separated by disease status of source patients. Error bars represent 95% confidence intervals determined using a bootstrapping approach.

## Discussion

In this study, we profile Treg heterogeneity within the lung tissue during infection with influenza A virus and introduce a new mouse model to track and deplete Treg subsets. The *Foxp3-iDTR* mouse model allows one to simultaneously track and delete distinct Treg subset. Previous systems to study Treg subsets have relied on deletion of specific floxed target genes within Tregs using *Foxp3-Cre* mice or deletion of *Foxp3* using *Foxp3*-floxed mice. While useful, these models are limited as they do not actually delete the Treg subset of interest but rather convert them into either a different type of Treg or an effector T-cell, respectively. Thus *Foxp3-iDTR* mice should be an invaluable tool for studying the roles of particular Treg subsets in a variety of tissues and disease settings. Using scRNA-Seq and our *Foxp3-iDTR* lineage tracking mice we demonstrate that in response to influenza A infection at least two transcriptionally distinct waves of Tregs populate the lungs: an initial ISG-Treg population that emerges at day 7 post-infection and contracts during clearance of influenza A, and a second wave of TR-Tregs that emerge during and following virus clearance. Importantly, the ISG-Treg and TR-Treg subsets have distinct TCR repertoires, suggesting that they recognize different antigens. The TR-Treg subset can be further broken down into a number of distinct transcriptional subsets, although these subsets have an overlapping TCR repertoire, suggesting that they recognize related antigens. Thus, the Treg response to influenza exhibits substantial diversity in kinetics of response, TCR repertoire, and transcriptome.

A number of key questions remain. First, it is unclear how the distinct Treg subsets observed during influenza A infection arise. A small subset of thymic (∼7%) and splenic Tregs (∼5%) expressing the ISG-signature can be found in the steady-state. These cells may be recruited to the lungs following viral infection. However, depletion of the vast majority of these pre-existing ISG-Tregs had no impact on the influx of ISG-Tregs to the lungs during a primary infection. It is unclear whether the ISG-Treg population in the lungs that arise after depletion of the pre-existing ISG-Treg repertoire comes from extremely rapid proliferation of the few ISG-Tregs that escape deletion, or possibly ISG-Tregs that fail to express sufficient *Mx1* to activate our reporter and thus are not deleted in our model. Alternatively, these could arise from Tregs that never encountered IFN previously, and hence did not express an ISG-signature initially, but acquired an ISG-signature during infection. Interestingly, a similar effect was not observed during a secondary infection with influenza, in which prior depletion of ISG-Tregs did result in a significant reduction in this subset of Tregs in the lung. One potential explanation is that prior to influenza infection only a fraction of the potential ISG-Treg pool acquires this gene signature during thymic development or at steady-state, but that after a primary infection all Tregs with this potential are labeled, allowing for their efficient depletion prior to a secondary infection.

A second question is what regulates the distinct temporal migration of ISG-Tregs and TR-Tregs. It is clear that these Treg subsets arrive in the lung with different kinetics but how this happens is not obvious. Likewise, where these Treg subsets localize within the lungs, and whether ISG-Tregs and TR-Tregs are found in different locations in the lungs, is also not clear.

Finally, the function of these distinct subsets of Tregs is unclear. For example, it remains unclear why distinct Treg subsets exist that uniquely express genes encoding endogenous opioids (*Penk*^+^), trefoil factor proteins (*Tff1*), or *Plac8*. A recent study demonstrated that activation of sensory neurons in the skin stimulated a Th17 response, indicative of crosstalk between the nervous and the immune systems (Cohen et al., 2019). PENK produced by Tregs may be acting on opioid receptors expressed by neurons in the lung to modulate pain perception. TFF1 is associated with mucus production in the gut (Järvå et al., 2020), although the function of TFF1-producing Tregs during IAV infection is unclear. The exact function of the ISG-Treg lineage also remains unclear although the observation that they correlate with improved response to COVID-19 suggests they play an important role. One possibility is that ISG-Tregs may be enriched for TCRs that recognize peptides derived from IFN stimulated genes, which are abundant during viral infections (Spencer et al., 2015). In this scenario, ISG-Tregs are drawn to an IFN-rich niche within the tissue and help to limit IFN-induced inflammation. In support of this possibility, recent studies have shown that limiting type I and type III IFN production following viral infections of the lung is required for tissue regeneration (Broggi et al., 2020; Major et al., 2020). Likewise, a recent study showed that depletion of Tregs during imiquimod-induced psoriasis led to uncontrolled IFNα and subsequent transcription of IFN-induced genes (Stockenhuber et al., 2018). Finally, a separate report demonstrated that house-dust mite (HDM)-reactive CD4^+^ T-cells and Tregs that express an IFN-signature are more frequent in asthmatic individuals without HDM allergy than those with HDM allergy (Seumois et al., 2020). Thus, ISG-Tregs are expanded in multiple inflammatory conditions. Identifying the antigens recognized by TR-Tregs and ISG-Tregs would likely provide key insights into their functional roles. Finally, in our analysis of Tregs in human patients with COVID-19, we identified a Treg subset expressing the cytokine IL32. As IL32 is not expressed in rodents (Kim, 2014) it was not identified in our murine scRNA-Seq studies. An important question is what causes the reciprocal relation between ISG-Tregs and *IL32*-Tregs in patients with mild versus severe COVID-19. The answer to that question could provide important insights into mechanisms leading to more severe disease in COVID-19 patients. Whether the distinct Treg subsets in the lung are functionally redundant or play non-overlapping roles during recovery from influenza A virus infection is also unclear. Using *Foxp3-DTR* mice we established that Tregs play an important role in limiting the frequency of IgA+ B cells in the lungs. Previous studies have shown that secreted IgA protects against influenza virus infection (van Riet et al., 2012). It is unclear why Tregs would limit this process, but they may play a role in ensuring preferential selection for high-affinity anti-viral B cells. Alternatively, limiting IgA^+^ B cells in the lung may be necessary to allow for proper repair of lung tissue post-infection. In conclusion, depletion of Tregs at the peak of the anti-influenza response does play an important role in subsequent immune function.

## Acknowledgements

We thank G. Hubbard, A. Rost, N. Keller, M. Dinh, T. Maiers, and J. Fiege for technical assistance, T. Martin, J. Motl and P. Champoux for cell sorting and maintenance of the Flow Cytometry Core Facility (UFCR), B. Ruis with the Genome Engineering Shared Resource (GESR) and Y. You with the Mouse Genetics Lab (MGL) for assistance developing the *Foxp3-iDTR* mice, the Minnesota Supercomputing Institute (MSI) at the University of Minnesota for providing computational resources. and Drs. Mukund and Subramaniam for providing the quality-filtered non-integrated human meta-analysis dataset. LES was supported by an immunology training grant (T32 AI997313) and an individual predoctoral F31 fellowship from the NIH (1F31AI154706-01A1). MAF was supported by NIH R01 AI124512 and R01 AI159554. This work was supported in part by NIH P30 CA77598 utilizing the following Masonic Cancer Center, University of Minnesota shared resource(s): UFCR, GESR and MGL.

## Author Contributions

Conceptualization, L.E.S., D.L.O. and M.A.F.; Formal analysis: T.P.K., C.H.O., B.M., and R.S. L.; Methodology, L.E.S.; Investigation, L.E.S., D.L.O. and S.J. Writing – Original Draft, L.E.S. and M.A.F..; Writing–Review & Editing, L.E.S., D.L.O., S.J., T.P.K., C.H.O., R.S.L., R.A.L. and M.A.F..; Funding Acquisition, L.E.S. and M.A.F.; Resources, R.A.L. and M.A.F.; Supervision, M.A.F.

## Data Availability Statement

All scRNA-Seq data is available at GEO under the following accession numbers: GSE123067, GSE195488, and GSE196909. Flow cytometry data is available upon request.

## Materials and Methods

### Mice

*Tclib* x *TCRα*^*+/-*^ x *Foxp3-GFP* (Jax stock #006772) mice were bred in house as previously described (Hsieh et al., 2004). *Tclib* x *TCRα*^*+/-*^ x *Foxp3-RFP* (Jax stock #008374) mice were bred in house. *Foxp3-iDTR* mice were created as described below and bred to *Mx1-Cre* (B6.Cg-Tg(Mx1-cre)1Cgn/J, Jax stock # 003556), *CD4-Cre* (B6.Cg-Tg(Cd4-cre)1Cwi/BfluJ, Jax stock # 022071) and *CD4-CreERT2* (B6(129X1)-Tg(Cd4-cre/ERT2)11Gnri/J, Jax stock #022356) mice.

### Foxp3-iDTR mice

A double-stranded DNA template was designed to contain a floxed stop cassette flanked by 800 base pair sequences homologous to the *Foxp3-DTR* locus. The DNA template, guide RNA, and CAS9 protein were micro-injected into *Foxp3-DTR* zygotes and zygotes were implanted into pseudopregnant female mice that had received prior injection of pregnant mare serum gonadotropin and human chorionic gonadotropin. Resulting pups were screened for the insertion using PCR primers 5’-CAAGTGCTCCAATCCCTGC-3’ and 5’-CTTGCATTCCTTTGGCGAGA-3’. Whole genome sequencing of tail DNA confirmed that the *Loxp-STOP-Loxp* cassette was integrated correctly and that there were no detectable off-target integrations or CAS9-induced mutations.

### Infections

Mice were anesthetized with a weight-determined dose of ketamine/xylazine and infected intranasally with 15 plaque-forming units (pfu) of influenza A virus (strain A/Puerto Rico/8/1934 H1N1). For depletion experiments, mice were administered 100ng of intraperitoneal DT on days 6, 7, and 8 post infection and 10ng of intranasal DT on day 7 post infection. Experiments involving mice were performed as dictated by the University of Minnesota Institutional Animal Care and Use Committee.

### IV antibody staining

IV antibody staining was performed as previously described (Anderson et al., 2014). Briefly, mice received an intravenous injection of 3ug of anti-CD4 (RM4-4, Biolegend 116022) antibody and were euthanized 3 minutes after injection.

### Isolation of lymphocytes

Mice were euthanized and the lymphoid tissues harvested into FACS buffer (1X PBS/2% FBS/0.05% sodium azide/2 mM EDTA, pH 7.4). Tissues were mashed between glass slides and filtered through 70um filter mesh to prepare single-cell suspensions. RBCs were lysed with ACK lysis buffer (150mM NH4Cl/10mM KHCO3/0.1mM Na2EDTA) for 5 minutes at room temperature.

### Isolation of lung lymphocytes

Lung lymphocytes were isolated as previously described (Fiege et al., 2019).

### Flow cytometry

Single-cell suspensions were washed with FACS buffer (1X PBS/2% FBS/2mM EDTA/0.05% sodium azide) and stained with ghost dye red 780 viability dye (Tonbo Biosciences 13-0865-T100) and antibodies against TCRβ (H57-597, BD Biosciences 742485), CD4 (RM4-5, Biolegend 100559), CD8 (53-6.7, Tonbo Biosciences 20-0081-U100), CD25 (PC61.5, Tonbo Biosciences 65-0251-U100), CD73 (TY/11.8, ThermoFisher Scientific 25-0731-82), B220 (RA3-6B2, BD Biosciences 563793), CD45 (30-F11, Biolegend 103155), CD90.2 (53.21 BD Biosciences 564365), GL7 (GL7, Biolegend 144604), and IgA (11-44-2, ThermoFisher Scientific 12-5994-81). Following surface staining, cells were fixed with the FOXP3 fixation and permeabilization kit and stained with antibodies against FOXP3 (FJK-16s ThermoFisher Scientific 17-5773-82, 48-5773-82), GFP (Rockland, 600-102-215), and IgG H+L (polyclonal, eBioscience 17-4012-82).

### Lung harvest, cell sorting, and cell hashtagging for single-cell RNA-seq

Mice received an intravenous injection of 3ug anti-CD4-biotin (RM4-4, Biolegend 116010) antibody 3 minutes prior to euthanasia. Lymphocytes from lungs were isolated into sort buffer (1XPBS/2% FBS/2mM EDTA) and processed as described above. Single-cell suspensions were stained with 2.5ug of anti-CD4-APC (RM4-5, Biolegend 100516) antibody in 1mL of sort buffer and incubated at 4°C for 30 minutes. Following wash, cells were resuspended in 50ul of anti-APC microbeads (Miltenyi) and 350ul of sort buffer. Cells were then washed and run over an LS column (Miltenyi). The bound fraction was collected and stained with ghost dye red 780 viability dye (Tonbo Biosciences 13-0865-T100), antibodies against TCRβ (H57-597, Biolegend 109229), CD8a (53-6.7, ThermoFisher Scientific 47-0081-82), Gr1 (RB6-8C5, ThermoFisher Scientific 47-5931-82), NK1.1 (PK136, ThermoFisher Scientific 47-5941-82), CD11b (M1/70, ThermoFisher Scientific 47-0112-82), CD11c (N418, ThermoFisher Scientific 47-0114-82), F4/80 (BM8, ThermoFisher Scientific 47-4801-82), and Ter119 (ThermoFisher Scientific 47-5921-82), 0.25ug total-seq C streptavidin-PE (C0951, Biolegend 405261), and 1ul total-seq C hashtag antibody (C0301-C0310, Biolegend) and incubated for 30 minutes at 4°C. Following wash, all samples were combined, resuspended in sort buffer, and run on the cell sorter.

### Single-cell RNA-sequencing of splenic Tregs

Spleens from *Tclib x TCRa*^*+/-*^ *x Foxp3-RFP* mice were dissociated as previously described (Owen et al., 2019) and stained with biotin-linked antibodies against CD8a and B220 as well as Ter119. After staining with streptavidin magnetic beads, cells were run over LS columns. The unbound fraction was used to sort CD4^+^CD8^−^ TCRVα2^+^FOXP3^+^ splenocytes into catch buffer. Antibodies against CD4 (GK1.5, BD Biosciences 563331), CD8a (53-6.7, ThermoFisher Scientific 47-0081-82), and TCRVα2 (B20.1, ThermoFisher Scientific 17-5812-82) were used. Cells were then centrifuged and resuspended in 1X PBS/10% FBS for single-cell capture. The 10X Genomics Chromium Next GEM Single Cell 5’ Kit (v2) was used to measure (1) mRNA transcript expression and (2) VDJ TCR repertoire. The library was sequenced on the NovaSeq 6000 with pair-end reads (26 bp for read 1 and 91 bp for read 2) (Illumina). Data available at accession number GSE195488.

### Single-cell RNA-sequencing of lung Tregs

To isolate Tregs, we sorted Dump^−^ TCRb^+^CD4^+^FOXP3-GFP^+^ cells into catch buffer (1XPBS/50% FBS). Cells were then centrifuged and resuspended in 1X PBS/10% FBS for single-cell capture. The 10X Genomics Chromium Next GEM Single Cell 5’ Kit (v2) was used to measure (1) mRNA transcript expression, (2) hashtag oligos (HTO), (3) CITE-seq oligos (ADT) for expression of cell-surface proteins and (4) VDJ TCR repertoire. To enhance TCRα chain CDR3 detection, given the fixed TCRβ allele, we only amplified the TCRα sequence using the following primer sets:

#### Mouse T cell Mix 1

Forward Primer (2uM final concentration): 5’-AATGATACGGCGACCACCGAGATCTACACTCTTTCCCTACACGACGCTC-3’

Reverse Outer Primer (0.5uM final concentration): 5’-CTGGTTGCTCCAGGCAATGG-3’

#### Mouse T cell Mix 2

Forward Primer (0.5uM final concentration): 5’-AATGATACGGCGACCACCGAGATCT-3’

Reverse Inner Primer (0.5uM final concentration): 5’-AGTCAAAGTCGGTGAACAGGCA-3’

Following library preparation, quality was assessed using a bioanalyzer and preliminary RNA-seq to determine approximate cell number and quality was performed on a MiSeq. After quality control, the libraries were sequenced on a NovaSeq 6000 with 2 × 150bp paired end reads. Raw and processed data are available through the GEO accession (GSE195488).

### Bioinformatics analysis of thymus and spleen single-cell RNA-sequencing data

The 10x Genomic Cell Ranger pipeline (version 3.0.0) was used to map reads to the mm10 reference (provided by 10X Genomics, version 3.0.0) and generate counts for each cell. Raw and processed data have been deposited at Gene Expression Omnibus and are available via GEO accession (see accession numbers above). Thymic and splenic T regulatory single cell expression data was analyzed from data initially collected in (Owen et al., 2019) (series GSE123067, dataset GSM3494565). Raw count data for both splenic and thymic single cell datasets were individually analyzed in R (version 3.5.0) using the Seurat package (version 3.1.1) and ggplot2 package (version 3.0.0). Each dataset was filtered for cells that expressed greater than 200 genes but less than 3200 genes. The remaining cells counts were normalized by a centered-log ratio method and principal component analysis (PCA) was performed on the 2000 most variable genes. Cells were clustered using the top 13 PC vectors for the splenic T cells and top 11 PC vectors for the thymic T cells using the FindNeighbors and FindClusters functions. Two-dimensional representations were generated using the RunTSNE and RunUMAP functions. Differential expression (wilcox) of clusters at different resolutions was performed to define biological significance. The interferon signature was determined to be best defined in the splenic T cell and thymic T cell cells at a resolution of 0.4 (cluster 3) and 0.35 (cluster 3) respectively.

### Bioinformatics analysis of lung Treg single-cell RNA-sequencing data

Raw sequencing data was processed using the 10X Genomics cellranger software (ver. 6.0.0) mkfastq function to demultiplex the Illumina libraries into gene expression (GEX), CITEseq expression (HTO and/or ADT), and T-cell receptor (TCR) datasets. The cellranger count function was used to align reads to the mouse reference genomes (refdata-gex-mm10-2020-A, refdata-cellranger-vdj-GRCm38-alts-ensembl-5.0.0; provided by 10X Genomics). See Tables 1 and 2 below. Raw count tables were loaded into R (ver. 4.0.3) and analyzed with the Seurat (ver. 4.0.1) or tidyverse (1.3.1) R packages. The GEX dataset was filtered to include only gelbeads in emulsion (GEMs, which are oil droplets containing uniquely barcoded beads that ideally contain one individual cell) expressing more than 300 genes (counts > 0) and genes expressed in more than 3 GEMs (counts > 0). The proportion of mitochondrial RNA in each GEM was calculated and GEMs with extreme levels (top 0.5% of all GEMs) were removed from the analysis. For the remaining GEMs, the raw HTO count table was supplied to GMM-demux software (ver. 0.2.1.3) (Xin et al., 2020) to classify which HTO tags were detected in each GEM. GEMs containing multiple HTOs (i.e. doublets or multiplets) were removed from downstream analysis. Initial analysis revealed no differences in HTO clustering between technical replicates and they were combined for all downstream analyses.

**Table 1:**
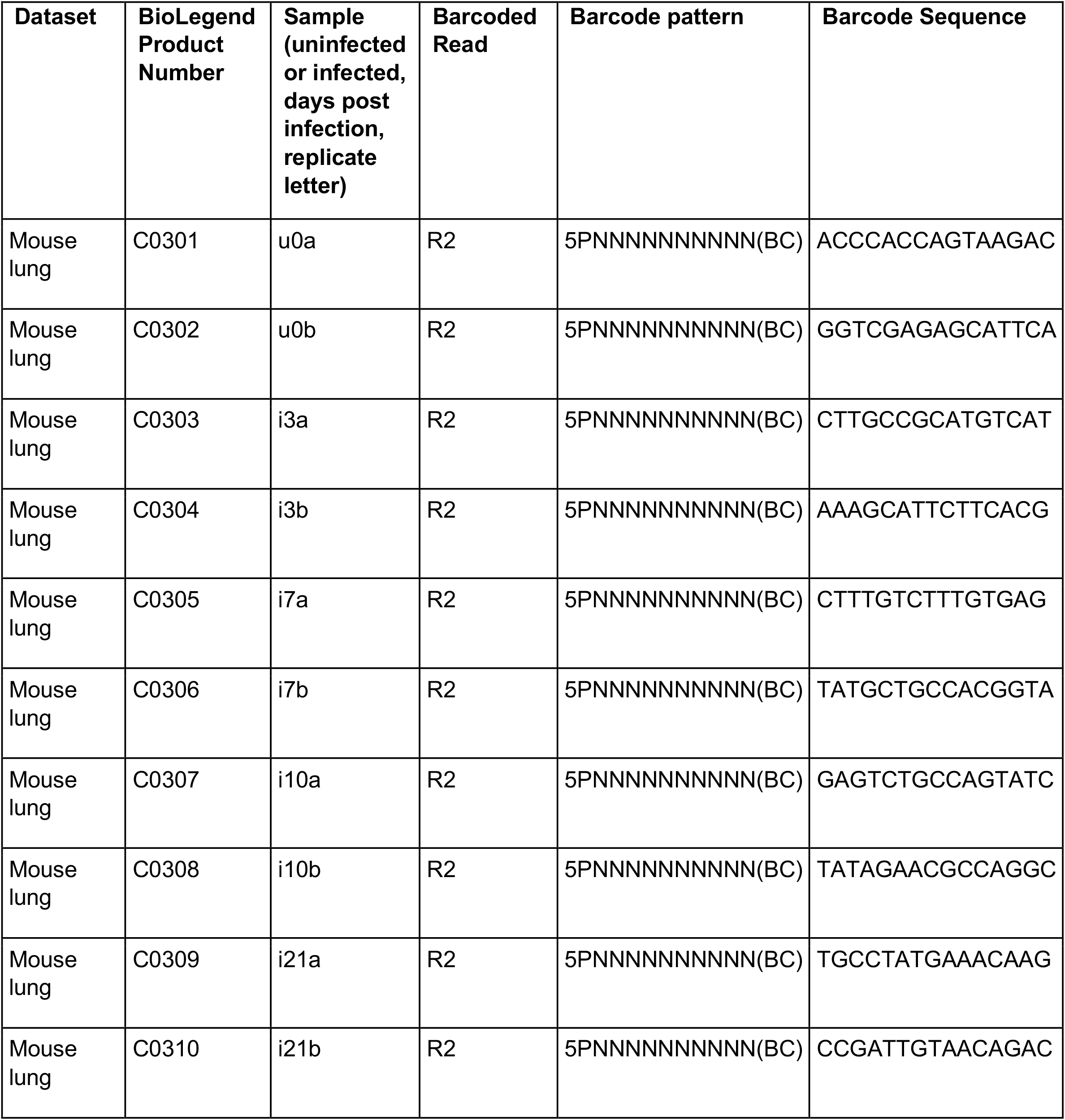
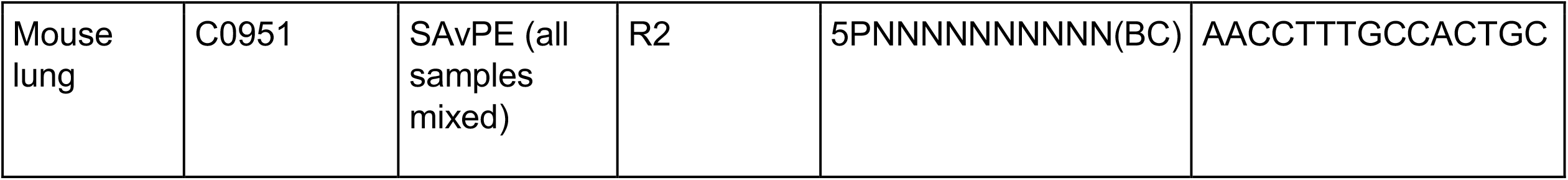
Hashtag (HTO) antibodies

**Table 2:**
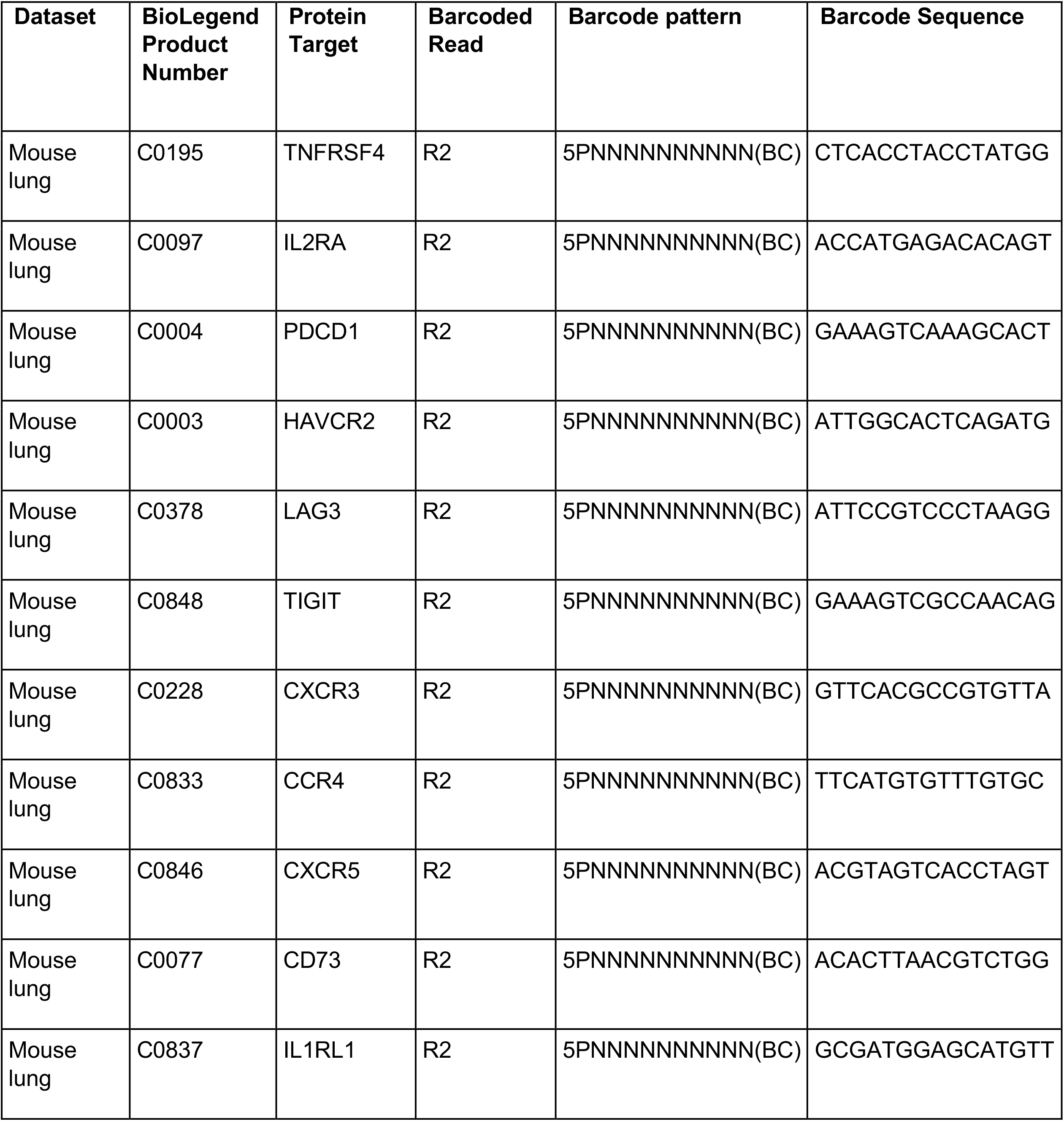
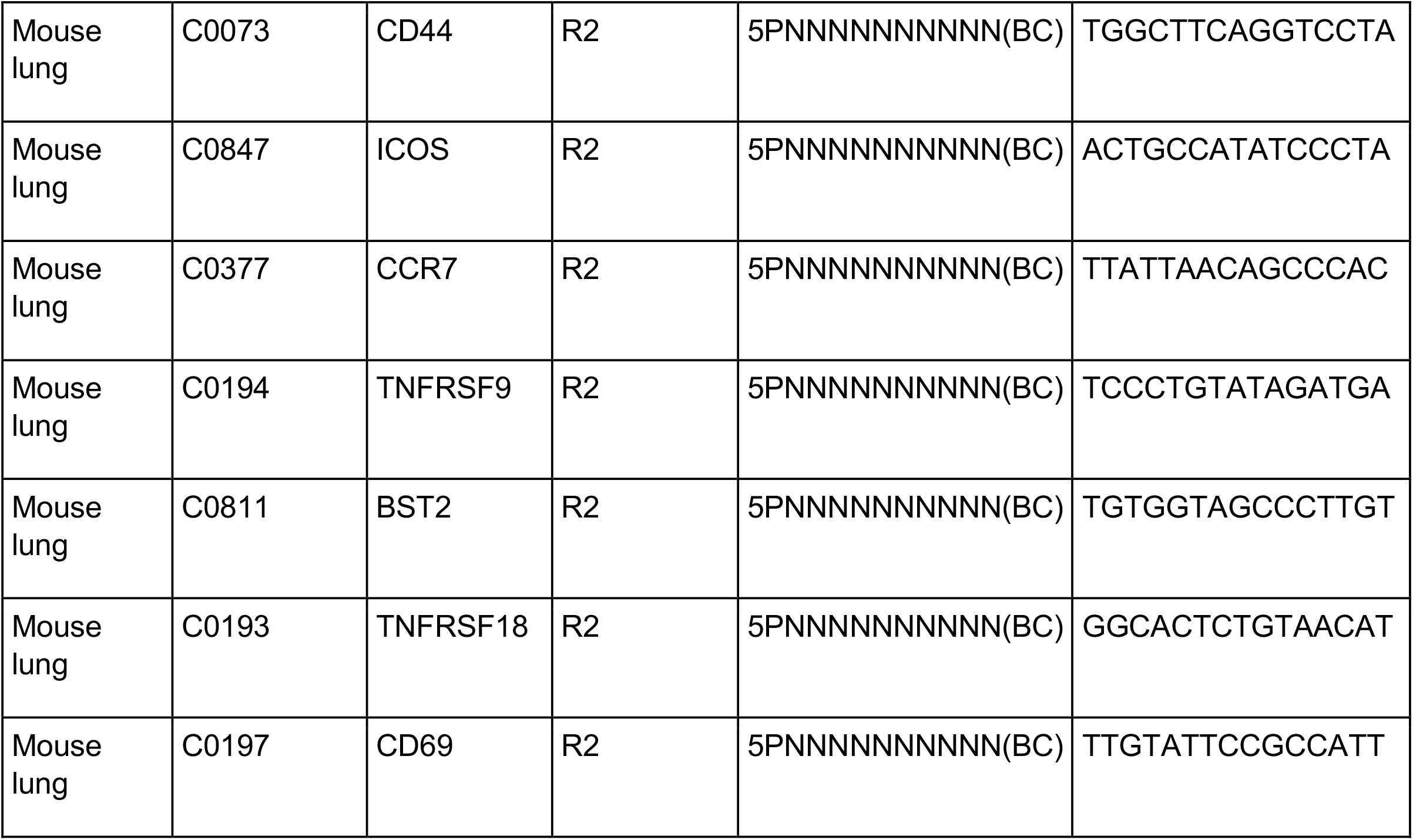
CITEseq (ADT) antibodies

Raw single cell surface protein expression data (ADT counts) were normalized according to the centered-log-ratio method using Seurat. Raw GEX counts were normalized and transformed using the Seurat SCTransform function (Hafemeister and Satija, 2019) including the percentage of mitochondria expression as a regression factor. Upon further analysis, we determined that four T-cell receptor genes (Trbv1, Trbv16, Trbv31 and Trav4-4-dv10) were driving the clustering of small unique clusters. The normalized gene expression levels for these genes were also used as regression factors in the final dataset. After regression the cells in these 4 TCR driven clusters clusters redistributed relatively uniformly throughout the UMAP. Each cell was classified according to its expression of canonical cell cycle genes using the Seurat CellCycleScoring function (S-phase and G2/M-phase gene sets provided by Seurat were originally developed by (Tirosh et al., 2016). Principal components analysis (PCA) was performed using the normalized, mean-centered, and scaled SCT dataset (RunPCA function). Two-dimensional projections were generated using the top 30 PCA vectors as input to RunUMAP function in Seurat. Cells were clustered using the FindNeighbors (top 30 PCA vectors) and FindClusters functions (testing a range of possible resolutions: 0.3, 0.4, 0.5, 0.6, 0.7). A final clustering resolution of 0.5 was selected.

Pairwise differential expression (DE) testing (Wilcox rank-sum) using the Seurat FindMarkers function was performed between all clusters. DE genes were significant based on log2-fold-change (≥ 0.25) and BH adjusted p-value (≤ 0.01). Figures were generated using the ggplot2 R package (ver. 2.3.3.3) (Wickham, 2016). The distribution of normalized GEX and ADT expression levels were displayed for cells/clusters using color heatmaps on UMAPs, dot plots, and tile plots.

A pseudotime cell trajectory analysis was completed using the Slingshot R package (ver. 1.8.0) (Street et al., 2018). The Seurat-based R object was converted for use with Slingshot using the Seurat function as.SingleCellExperiment. A trajectory was inferred with the slingshot function using cluster labels, UMAP coordinates, and the central Tregs (cluster 1) as the root of the trajectory. Cell groups were compared (e.g. uninfected vs. infected) along each of the slingshot lineages using a differential topology approach. To determine statistical significance, a permutation test was performed. For each cell group, the weighted mean of pseudotime values was calculated using the slingshot curve weights. An initial test-statistic was calculated as the difference in weighted means between the two cell groups. A null distribution was generated by randomization of cell group labels and resampling 10,000 times. P-values were calculated as the number of test-statistics generated under the null distribution that were as or more extreme than the initial test-statistic (i.e., using real cell group labels).

Raw and processed data have been deposited at the Gene Expression Omnibus (GEO) database (accession number: GSE196909).

### Whole Lung RNA-seq

Whole lung was added to 3mL of buffer RLT with β-mercaptoethanol and reagent DX and digested in M-tubes on the GentleMacs RNA setting. After a quick spin to remove debris, lung homogenate was aliquoted and 300ul was processed through the Qiagen RNeasy Mini Kit to extract total RNA. The RNA-seq library was prepared using the TruSeq Stranded kit. Samples were sequenced on the NovaSeq 6000 using 150 base-pair paired-end reads for a total of 20 million reads per sample.

Bulk RNAseq libraries were run through the CHURP pipeline as previously described (https://dl.acm.org/doi/10.1145/3332186.3333156). Briefly, this wrapper quality trims reads with Trimmomatic (Bolger et al., 2014), aligns to the human reference genome (GRCh38) using HISAT2 (Kim et al., 2019), and converts alignments to feature counts using SAMTools (Li et al., 2009) and featureCounts() from the Rsubread package (Liao et al., 2019). Expression of genes present in the Ensembl GRCh38 v100 annotation set and longer than 300 bases were further quantified using edgeR (Robinson et al., 2010). Features present in two or more samples at more than 1 count per million were selected to calculate normalization factors with method “RLE” using calcNormFactors(). Dispersion was then estimated using estimateDisp() and differentially expressed genes were identified using glmQLFTest() with default parameters. Volcano plots were generated using ggplot2 (Wickham, 2016).

GSEA analyses were performed on genes lists consisting of genes with at least a 2-fold expression difference and a false-discovery corrected p-value less than 0.1. Genes were rank ordered based direction of difference followed by the FDR-corrected p-value of the comparison. These gene lists were analyzed using the R package clusterProfiler (Wu et al., 2021) and compared against human HALLMARK and GO term gene lists using an q-value cut off of 0.1. Dotplots were generated through ClusterProfiler using the dotplot() function from the R package enrichplot (Wu et al., 2021).

### Bioinformatics analysis of human COVID-19 dataset

The dataset described in section 4.3 Mukund et al. (2021) was made available to us by the paper’s authors. The dataset was already filtered for the criteria described in section 4.3 but not integrated. Using available metadata, the dataset was separated into the 5 original datasets and processed and integrated using Seurat’s (v. 4.0.5) IntegrateData function. The same parameters were used as described in section 4.3 of Mukund et al 2021. Following integration, cells annotated by Mukund et al. (Mukund et al., 2021) as CD4 or CD8 T-cells were subsampled from the original dataset and processed as described in section 4.4 of Mukund et al., 2021, and re-clustered at a range of resolutions (0.8 – 1.2). A resolution value of 0.8 was used and cluster 9 (CD4^+^FOXP3^+^CD25^+^IL7R^lo^ T cells) was subsampled for further analysis. The CD4^+^FOXP3^+^CD25^+^IL7R^lo^ cell subset was processed using the same parameters described in section 4.4 of Mukund et al., 2021. Clustering was performed at a range of resolutions (0.1 – 1) and the clusters identified at a resolution value of 0.3 were used in further analyses. Confidence bars were calculated for the resolution 0.3 clusters using single cell differential composition (scDC) analysis (v. 0.1.0) (Cao et al., 2019). The scDC_noClustering function with the bias corrected and accelerated (BCa) confidence interval calculation method was used. All work was done in R (v. 4.1.2).

### Statistics

In the graphs horizontal bars represents means and vertical bars represent standard deviations. In the text, +/- values represent standard deviations. Normality was assessed using Shapiro-Wilks test. For comparisons of two groups, an unpaired t-test was used for normal data and Mann-Whitney test was used for non-normal data. For comparisons of two groups across multiple time points, correction for multiple comparisons was performed using the Holm-Sidak method. A two-way ANOVA was used to compare three or more groups containing two factors.

---

**Supplemental Figure 1.**
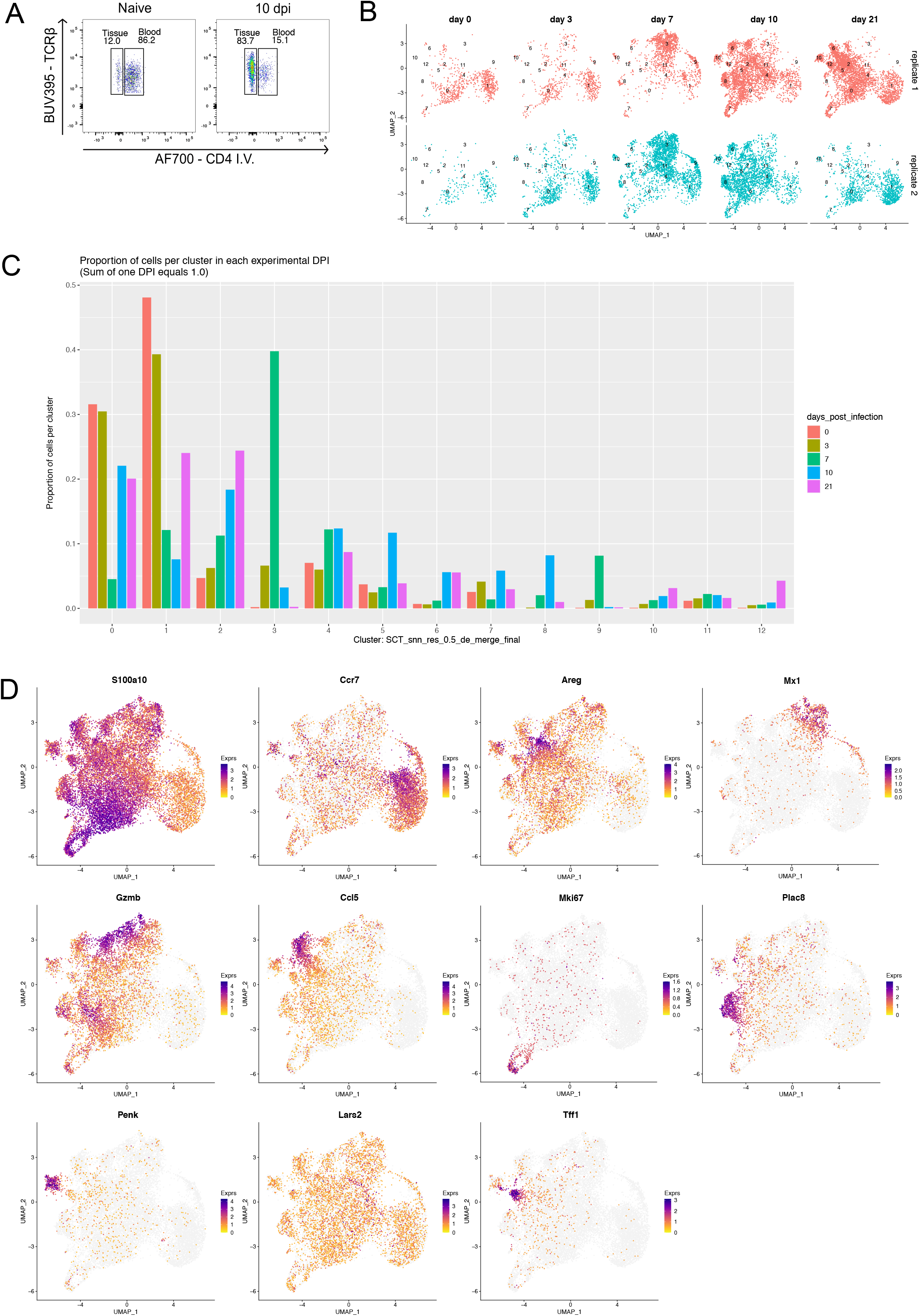
(A) IV-labeling of lung Tregs in naive mice and mice at 10 days post influenza A infection. Cells gated on live>TCRβ+>CD4+>Foxp3+. (B) UMAPs of replicate mice at each time point post infection (C) Frequency of Tregs from each time point that are within each cluster. (D) Feature plots of defining genes for each cluster.

**Supplemental Figure 2.**
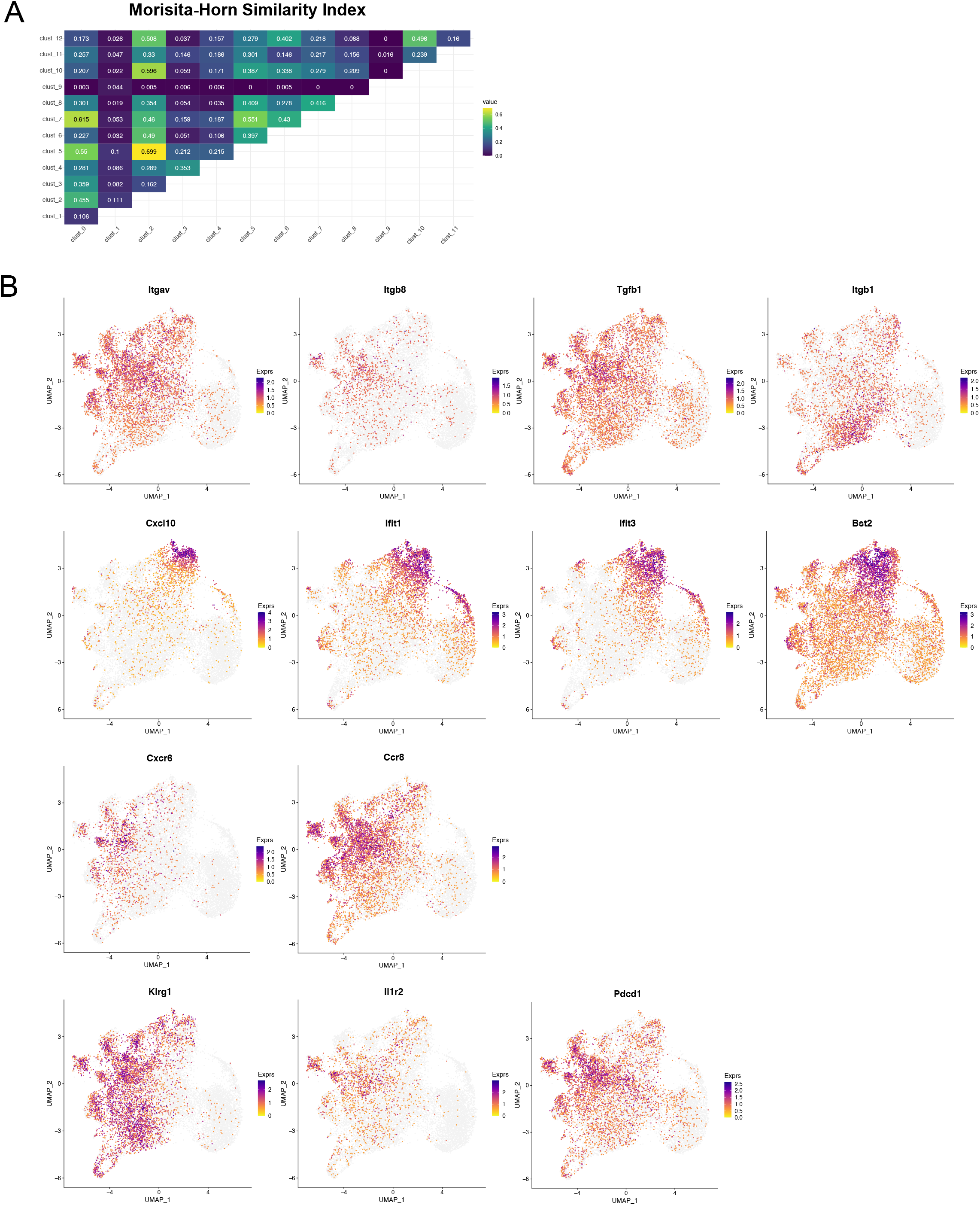
(A) Morisita-horn similarity index of TCRs from each UMAP cluster. (B) Feature plots of transcripts defining IFNsig-Treg cluster and tissue-repair Treg clusters.

**Supplemental Figure 3.**
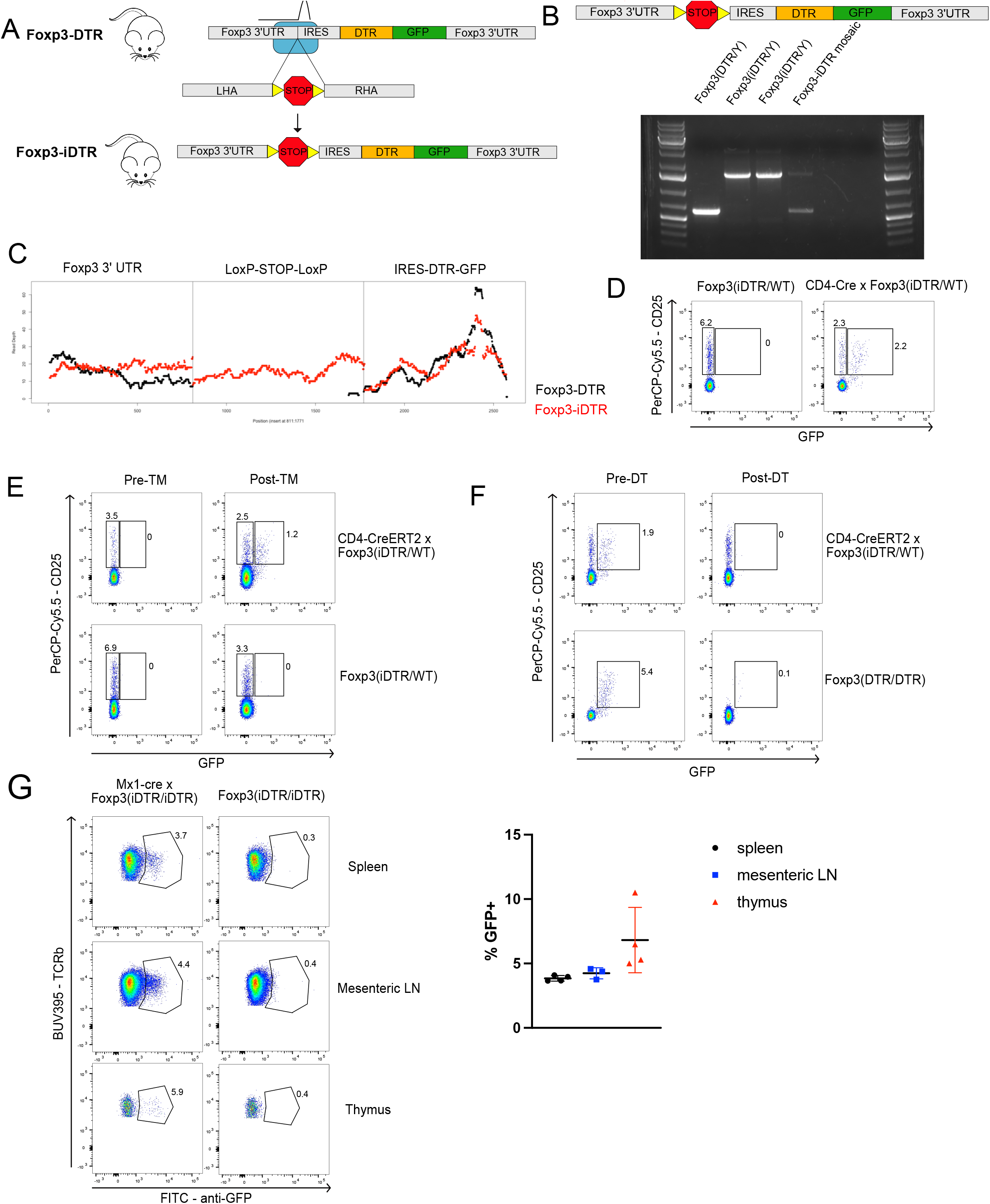
(A) Model of Fo*xp*3-*i*DTR mouse. (B) PCR amplification of the floxed-STOP cassette in the Foxp3-iDTR mice. Red arrows represent PCR primers. (C) Whole genome sequencing reads from the Foxp3-DTR and Foxp3-iDTR mice. (D) Expression of GFP in CD4-Cre x Foxp3-iDTR mice. Representative of 6 mice. (E) Expression of GFP in CD4-CreERT2 x Foxp3-iDTR mice pre- and post-tamoxifen treatment. Representative of 7 mice. (F) Expression of GFP pre- and post-DT treatment. Representative of 4 mice. (G) Frequency of ISG-Tregs in the spleen, mesenteric LN, and thymus of Mx1-Cre x Foxp3-iDTR mice at the steady state. The horizontal bars represent the mean and vertical bars represent the standard deviation.

**Supplemental Figure 4.**
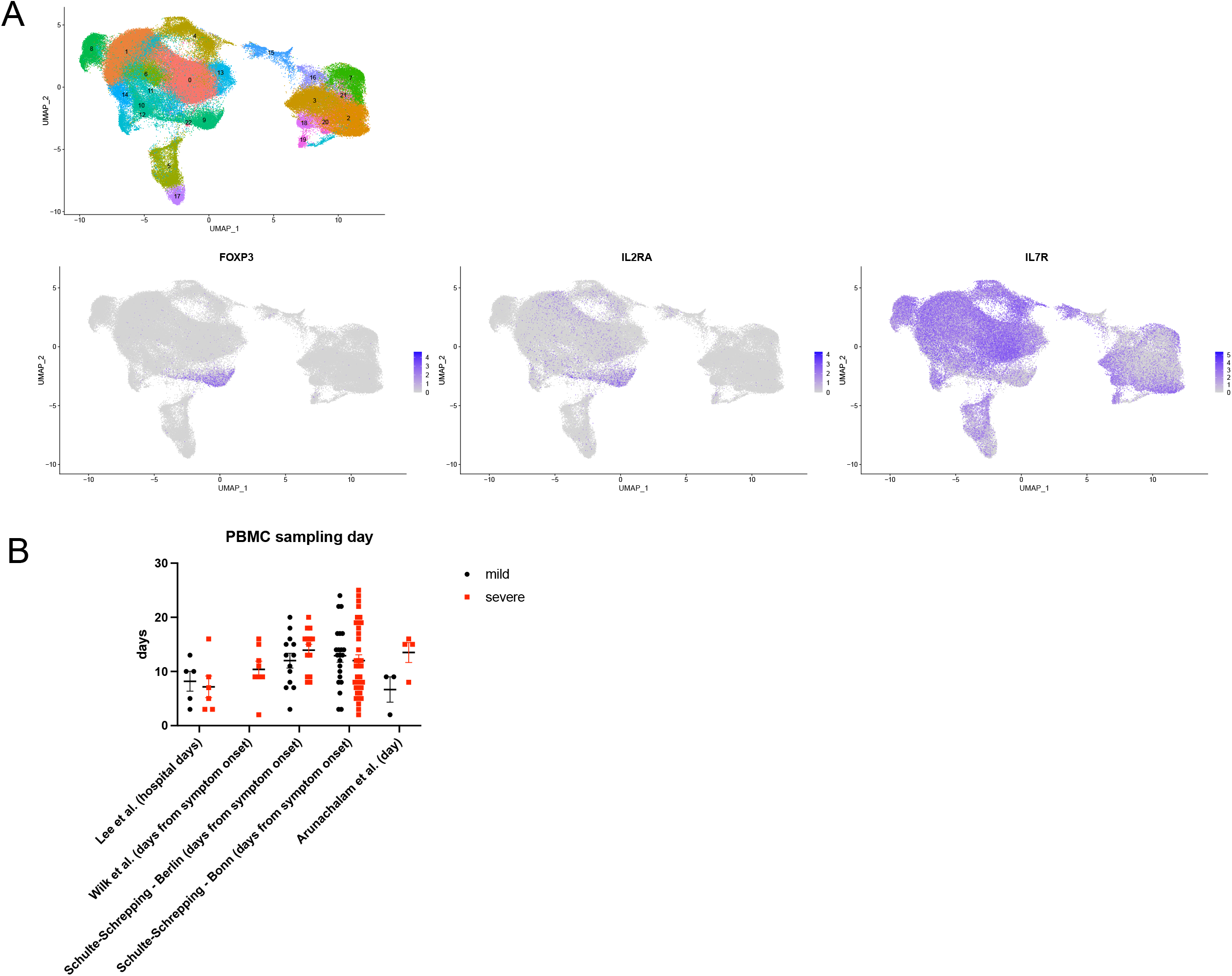
(A) UMAP of PBMCs from healthy people and COVID-19 patients. Feature plots of FOXP3, IL2RA, and IL7R. (B) Sampling day of mild and severe patients’ PBMCs in each study of *FOXP3, IL2RA*, and *IL7R*. (B) Sampling day of mild and severe patients’ PBMCs in each study included in meta-analysis. Horizontal bars represent means and vertical bars represent standard deviations. 2-way ANOVA showed that there was no statistical difference between the sampling days of mild and severe patients (p=0.7304) or the sampling days of the studies included in the meta-analysis (p=0.0660). For mild patients, n=43. and for severe patients, n=68.

## References

Anderson, K.G., K. Mayer-Barber, H. Sung, L. Beura, B.R. James, J.J. Taylor, L. Qunaj, T.S. Griffith, V. Vezys, D.L. Barber, and D. Masopust. 2014. Intravascular staining for discrimination of vascular and tissue leukocytes. Nat Protoc 9:209–222.

Arpaia, N., J.A. Green, B. Moltedo, A. Arvey, S. Hemmers, S. Yuan, P.M. Treuting, and A.Y. Rudensky. 2015. A Distinct Function of Regulatory T Cells in Tissue Protection. Cell 162:1078–1089.

Broggi, A., S. Ghosh, B. Sposito, R. Spreafico, F. Balzarini, A. Lo Cascio, N. Clementi, M. De Santis, N. Mancini, F. Granucci, and I. Zanoni. 2020. Type III interferons disrupt the lung epithelial barrier upon viral recognition. Science 369:706–712.

Chaudhry, A., D. Rudra, P. Treuting, R.M. Samstein, Y. Liang, A. Kas, and A.Y. Rudensky. 2009. CD4+ regulatory T cells control TH17 responses in a Stat3-dependent manner. Science 326:986–991.

Cohen, J.A., T.N. Edwards, A.W. Liu, T. Hirai, M.R. Jones, J. Wu, Y. Li, S. Zhang, J. Ho, B.M. Davis, K.M. Albers, and D.H. Kaplan. 2019. Cutaneous TRPV1. Cell 178:919-932.e914.

Delacher, M., C.D. Imbusch, A. Hotz-Wagenblatt, J.P. Mallm, K. Bauer, M. Simon, D. Riegel, A.F. Rendeiro, S. Bittner, L. Sanderink, A. Pant, L. Schmidleithner, K.L. Braband, B. Echtenachter, A. Fischer, V. Giunchiglia, P. Hoffmann, M. Edinger, C. Bock, M. Rehli, B. Brors, C. Schmidl, and M. Feuerer. 2020. Precursors for Nonlymphoid-Tissue Treg Cells Reside in Secondary Lymphoid Organs and Are Programmed by the Transcription Factor BATF. Immunity 52:295-312.e211.

Harrison, O.J., J.L. Linehan, H.Y. Shih, N. Bouladoux, S.J. Han, M. Smelkinson, S.K. Sen, A.L. Byrd, M. Enamorado, C. Yao, S. Tamoutounour, F. Van Laethem, C. Hurabielle, N. Collins, A. Paun, R. Salcedo, J.J. O’Shea, and Y. Belkaid. 2019. Commensal-specific T cell plasticity promotes rapid tissue adaptation to injury. Science 363:

Josefowicz, S.Z., L.F. Lu, and A.Y. Rudensky. 2012. Regulatory T cells: mechanisms of differentiation and function. Annual review of immunology 30:531–564.

Järvå, M.A., J.P. Lingford, A. John, N.M. Soler, N.E. Scott, and E.D. Goddard-Borger. 2020. Trefoil factors share a lectin activity that defines their role in mucus. Nat Commun 11:2265.

Kim, J.M., J.P. Rasmussen, and A.Y. Rudensky. 2007. Regulatory T cells prevent catastrophic autoimmunity throughout the lifespan of mice. Nature immunology 8:191–197.

Kim, S. 2014. Interleukin-32 in inflammatory autoimmune diseases. Immune Netw 14:123–127.

Koch, M.A., G. Tucker-Heard, N.R. Perdue, J.R. Killebrew, K.B. Urdahl, and D.J. Campbell. 2009. The transcription factor T-bet controls regulatory T cell homeostasis and function during type 1 inflammation. Nat Immunol 10:595–602.

Kühn, R., F. Schwenk, M. Aguet, and K. Rajewsky. 1995. Inducible gene targeting in mice. Science 269:1427–1429.

Lee, J.S., and E.C. Shin. 2020. The type I interferon response in COVID-19: implications for treatment. Nat Rev Immunol 20:585–586.

Levine, A.G., A. Mendoza, S. Hemmers, B. Moltedo, R.E. Niec, M. Schizas, B.E. Hoyos, E.V. Putintseva, A. Chaudhry, S. Dikiy, S. Fujisawa, D.M. Chudakov, P.M. Treuting, and A.Y. Rudensky. 2017. Stability and function of regulatory T cells expressing the transcription factor T-bet. Nature 546:421–425.

Loebbermann, J., H. Thornton, L. Durant, T. Sparwasser, K.E. Webster, J. Sprent, F.J. Culley, C. Johansson, and P.J. Openshaw. 2012. Regulatory T cells expressing granzyme B play a critical role in controlling lung inflammation during acute viral infection. Mucosal Immunol 5:161–172.

Lu, D.R., H. Wu, I. Driver, S. Ingersoll, S. Sohn, S. Wang, C.M. Li, and H. Phee. 2020. Dynamic changes in the regulatory T-cell heterogeneity and function by murine IL-2 mutein. Life Sci Alliance 3:

Major, J., S. Crotta, M. Llorian, T.M. McCabe, H.H. Gad, S.L. Priestnall, R. Hartmann, and A. Wack. 2020. Type I and III interferons disrupt lung epithelial repair during recovery from viral infection. Science 369:712–717.

Miragaia, R.J., T. Gomes, A. Chomka, L. Jardine, A. Riedel, A.N. Hegazy, N. Whibley, A. Tucci, X. Chen, I. Lindeman, G. Emerton, T. Krausgruber, J. Shields, M. Haniffa, F. Powrie, and S.A. Teichmann. 2019. Single-Cell Transcriptomics of Regulatory T Cells Reveals Trajectories of Tissue Adaptation. Immunity 50:493-504.e497.

Mukund, K., P. Nayak, C. Ashokkumar, S. Rao, J. Almeda, M.M. Betancourt-Garcia, R. Sindhi, and S. Subramaniam. 2021. Immune Response in Severe and Non-Severe Coronavirus Disease 2019 (COVID-19) Infection: A Mechanistic Landscape. Frontiers in immunology 12:738073.

Owen, D.L., R.S. La Rue, S.A. Munro, and M.A. Farrar. 2022. Tracking Regulatory T Cell Development in the Thymus Using scRNA-Seq/TCR-Seq. J Immunol

Owen, D.L., S.A. Mahmud, L.E. Sjaastad, J.B. Williams, J.A. Spanier, D.R. Simeonov, R. Ruscher, W. Huang Proekt, C.N. Miller, C. Hekim, J.C. Jeschke, P. Aggarwal, U. Broeckel, R.S. LaRue, C.M. Henzler, M.L. Alegre, M.S. Anderson, A. August, A. Marson, Y. Zheng, C.B. Williams, and M.A. Farrar. 2019. Thymic regulatory T cells arise via two distinct developmental programs. Nature immunology 20:195–205.

Peligero-Cruz, C., T. Givony, A. Sebé-Pedrós, J. Dobeš, N. Kadouri, S. Nevo, F. Roncato, R. Alon, Y. Goldfarb, and J. Abramson. 2020. IL18 signaling promotes homing of mature Tregs into the thymus. Elife 9:

Permanyer, M., B. Bošnjak, S. Glage, M. Friedrichsen, S. Floess, J. Huehn, G.E. Patzer, I. Odak, N. Eckert, R. Zargari, L. Ospina-Quintero, H. Georgiev, and R. Förster. 2021. Efficient IL-2R signaling differentially affects the stability, function, and composition of the regulatory T-cell pool. Cell Mol Immunol 18:398–414.

Playford, R.J., T. Marchbank, R.A. Goodlad, R.A. Chinery, R. Poulsom, and A.M. Hanby. 1996. Transgenic mice that overexpress the human trefoil peptide pS2 have an increased resistance to intestinal damage. Proc Natl Acad Sci U S A 93:2137–2142.

Sefik, E., N. Geva-Zatorsky, S. Oh, L. Konnikova, D. Zemmour, A.M. McGuire, D. Burzyn, A. Ortiz-Lopez, M. Lobera, J. Yang, S. Ghosh, A. Earl, S.B. Snapper, R. Jupp, D. Kasper, D. Mathis, and C. Benoist. 2015. MUCOSAL IMMUNOLOGY. Individual intestinal symbionts induce a distinct population of RORγ⁺ regulatory T cells. Science 349:993–997.

Seumois, G., C. Ramirez-Suastegui, B.J. Schmiedel, S. Liang, B. Peters, A. Sette, and P. Vijayanand. 2020. Single-cell transcriptomic analysis of allergen-specific T cells in allergy and asthma. Sci Immunol 5:

Shime, H., M. Odanaka, M. Tsuiji, T. Matoba, M. Imai, Y. Yasumizu, R. Uraki, K. Minohara, M. Watanabe, A.J. Bonito, H. Fukuyama, N. Ohkura, S. Sakaguchi, A. Morita, and S. Yamazaki. 2020. Proenkephalin. Proc Natl Acad Sci U S A 117:20696–20705.

Sjaastad, L.E., D.L. Owen, S.I. Tracy, and M.A. Farrar. 2021. Phenotypic and Functional Diversity in Regulatory T Cells. Front Cell Dev Biol 9:715901.

Smigiel, K.S., E. Richards, S. Srivastava, K.R. Thomas, J.C. Dudda, K.D. Klonowski, and D.J. Campbell. 2014. CCR7 provides localized access to IL-2 and defines homeostatically distinct regulatory T cell subsets. J Exp Med 211:121–136.

Spencer, C.T., J.S. Bezbradica, M.G. Ramos, C.D. Arico, S.B. Conant, P. Gilchuk, J.J. Gray, M. Zheng, X. Niu, W. Hildebrand, A.J. Link, and S. Joyce. 2015. Viral infection causes a shift in the self peptide repertoire presented by human MHC class I molecules. Proteomics Clin Appl 9:1035–1052.

Stockenhuber, K., A.N. Hegazy, N.R. West, N.E. Ilott, A. Stockenhuber, S.J. Bullers, E.E. Thornton, I.C. Arnold, A. Tucci, H. Waldmann, G.S. Ogg, and F. Powrie. 2018. Foxp3. J Exp Med 215:1987–1998.

van Riet, E., A. Ainai, T. Suzuki, and H. Hasegawa. 2012. Mucosal IgA responses in influenza virus infections; thoughts for vaccine design. Vaccine 30:5893–5900.

Veiga-Parga, T., S. Sehrawat, and B.T. Rouse. 2013. Role of regulatory T cells during virus infection. Immunol Rev 255:182–196.

Wohlfert, E.A., J.R. Grainger, N. Bouladoux, J.E. Konkel, G. Oldenhove, C.H. Ribeiro, J.A. Hall, R. Yagi, S. Naik, R. Bhairavabhotla, W.E. Paul, R. Bosselut, G. Wei, K. Zhao, M. Oukka, J. Zhu, and Y. Belkaid. 2011. GATA3 controls Foxp3⁺ regulatory T cell fate during inflammation in mice. J Clin Invest 121:4503–4515.

Worthington, J.J., A. Kelly, C. Smedley, D. Bauché, S. Campbell, J.C. Marie, and M.A. Travis. 2015. Integrin αvβ8-Mediated TGF-β Activation by Effector Regulatory T Cells Is Essential for Suppression of T-Cell-Mediated Inflammation. Immunity 42:903–915.

Zemmour, D., R. Zilionis, E. Kiner, A.M. Klein, D. Mathis, and C. Benoist. 2018. Single-cell gene expression reveals a landscape of regulatory T cell phenotypes shaped by the TCR. Nat Immunol 19:291–301.

Zheng, Y., A. Chaudhry, A. Kas, P. deRoos, J.M. Kim, T.T. Chu, L. Corcoran, P. Treuting, U. Klein, and A.Y. Rudensky. 2009. Regulatory T-cell suppressor program co-opts transcription factor IRF4 to control T(H)2 responses. Nature 458:351–356.

## References

Bolger, A.M., M. Lohse, and B. Usadel. 2014. Trimmomatic: a flexible trimmer for Illumina sequence data. Bioinformatics 30:2114–2120.

Cao, Y., Y. Lin, J.T. Ormerod, P. Yang, J.Y.H. Yang, and K.K. Lo. 2019. scDC: single cell differential composition analysis. BMC Bioinformatics 20:721.

Fiege, J.K., I.A. Stone, R.E. Dumm, B.M. Waring, B.T. Fife, J. Agudo, B.D. Brown, N.S. Heaton, and R.A. Langlois. 2019. Long-term surviving influenza infected cells evade CD8+ T cell mediated clearance. PLoS Pathog 15:e1008077.

Hafemeister, C., and R. Satija. 2019. Normalization and variance stabilization of single-cell RNA-seq data using regularized negative binomial regression. Genome Biol 20:296.

Kim, D., J.M. Paggi, C. Park, C. Bennett, and S.L. Salzberg. 2019. Graph-based genome alignment and genotyping with HISAT2 and HISAT-genotype. Nat Biotechnol 37:907–915.

Li, H., B. Handsaker, A. Wysoker, T. Fennell, J. Ruan, N. Homer, G. Marth, G. Abecasis, R. Durbin, and G.P.D.P. Subgroup. 2009. The Sequence Alignment/Map format and SAMtools. Bioinformatics 25:2078–2079.

Liao, Y., G.K. Smyth, and W. Shi. 2019. The R package Rsubread is easier, faster, cheaper and better for alignment and quantification of RNA sequencing reads. Nucleic Acids Res 47:e47.

Mukund, K., P. Nayak, C. Ashokkumar, S. Rao, J. Almeda, M.M. Betancourt-Garcia, R. Sindhi, and S. Subramaniam. 2021. Immune Response in Severe and Non-Severe Coronavirus Disease 2019 (COVID-19) Infection: A Mechanistic Landscape. Front Immunol 12:738073.

Owen, D.L., S.A. Mahmud, L.E. Sjaastad, J.B. Williams, J.A. Spanier, D.R. Simeonov, R. Ruscher, W. Huang, I. Proekt, C.N. Miller, C. Hekim, J.C. Jeschke, P. Aggarwal, U. Broeckel, R.S. LaRue, C.M. Henzler, M.L. Alegre, M.S. Anderson, A. August, A. Marson, Y. Zheng, C.B. Williams, and M.A. Farrar. 2019. Thymic regulatory T cells arise via two distinct developmental programs. Nat Immunol 20:195–205.

Robinson, M.D., D.J. McCarthy, and G.K. Smyth. 2010. edgeR: a Bioconductor package for differential expression analysis of digital gene expression data. Bioinformatics 26:139–140.

Street, K., D. Risso, R.B. Fletcher, D. Das, J. Ngai, N. Yosef, E. Purdom, and S. Dudoit. 2018. Slingshot: cell lineage and pseudotime inference for single-cell transcriptomics. BMC Genomics 19:477.

Tirosh, I., B. Izar, S.M. Prakadan, M.H. Wadsworth, D. Treacy, J.J. Trombetta, A. Rotem, C. Rodman, C. Lian, G. Murphy, M. Fallahi-Sichani, K. Dutton-Regester, J.R. Lin, O. Cohen, P. Shah, D. Lu, A.S. Genshaft, T.K. Hughes, C.G. Ziegler, S.W. Kazer, A. Gaillard, K.E. Kolb, A.C. Villani, C.M. Johannessen, A.Y. Andreev, E.M. Van Allen, M. Bertagnolli, P.K. Sorger, R.J. Sullivan, K.T. Flaherty, D.T. Frederick, J. Jané-Valbuena, C.H. Yoon, O. Rozenblatt-Rosen, A.K. Shalek, A. Regev, and L.A. Garraway. 2016. Dissecting the multicellular ecosystem of metastatic melanoma by single-cell RNA-seq. Science 352:189–196.

Wickham, H. 2016. ggplot2: Elegant Graphics for Data Analysis. Springer-Verlag New York,

Wu, T., E. Hu, S. Xu, M. Chen, P. Guo, Z. Dai, T. Feng, L. Zhou, W. Tang, L. Zhan, X. Fu, S. Liu, X. Bo, and G. Yu. 2021. clusterProfiler 4.0: A universal enrichment tool for interpreting omics data. Innovation (Camb) 2:100141.

Xin, H., Q. Lian, Y. Jiang, J. Luo, X. Wang, C. Erb, Z. Xu, X. Zhang, E. Heidrich-O’Hare, Q. Yan, R.H. Duerr, K. Chen, and W. Chen. 2020. GMM-Demux: sample demultiplexing, multiplet detection, experiment planning, and novel cell-type verification in single cell sequencing. Genome Biol 21:188.

